# Functional brain adaptations during speech processing in 4-month-old bilingual infants

**DOI:** 10.1101/2024.01.26.577342

**Authors:** Borja Blanco, Monika Molnar, Irene Arrieta, César Caballero-Gaudes, Manuel Carreiras

## Abstract

Language learning is influenced by both neural development and environmental experiences. This work investigates the influence of early bilingual experience on the neural mechanisms underlying speech processing in 4-month-old infants. We study how an early environmental factor such as bilingualism interacts with neural development by comparing monolingual and bilingual infants’ brain responses to speech. We used functional near-infrared spectroscopy (fNIRS) to measure 4-month-old Spanish-Basque bilingual and Spanish monolingual infants’ brain responses while they listened to forward (FW) and backward (BW) speech stimuli in Spanish. We reveal distinct neural signatures associated with bilingual adaptations, including increased engagement of bilateral inferior frontal and temporal regions during speech processing in bilingual infants, as opposed to left hemispheric specialization observed in monolingual infants. This study provides compelling evidence of bilingualism-induced brain adaptations during speech processing in infants as young as four months. These findings emphasize the role of early language experience in shaping neural plasticity during infancy suggesting that bilingual exposure at this young age profoundly influences the neural mechanisms underlying speech processing.

**Research Highlights:**

- This research sheds light on the intricate relationship between language learning, neural maturation, and environmental factors in early infancy.
- Bilingual infants exhibit unique functional brain adaptations during speech processing as early as 4 months of age.
- This study underscores the critical role of language experience in shaping the developing brain.
- These findings have implications for our understanding of language acquisition and neuroplasticity in infants growing up in bilingual environments.

## Introduction

Language learning is mediated by both neural maturation and experience with one’s environment. Here, we focus on how environmental factors interplay with neural development. Specifically, we look at how the type of exposure to spoken language during the first months of life shapes the developing human brain. To do this, we compare the neural responses of monolingual and bilingual infants to speech. While the amount of language exposure should be comparable across monolingual and bilingual environments, bilingual infants likely receive less exposure to each of their languages compared to their monolingual peers, because their exposure time is split between two inputs. In addition, bilingual infants need to perform specific computations, such as separating their languages or storing the information of two linguistic inputs (e.g., Costa & Sebastián-Gallés, 2014).

Previous research demonstrated that a bilingual context might modulate infants’ behavioural responses to familiar vs. unfamiliar languages. For example, English monolingual newborns prefer English over Tagalog, while English-Tagalog bilingual newborns respond similarly to both native languages (Byers-Heinlein et al., 2010; Mehler et al., 1988; Moon et al., 1993). Early bilingual exposure also leads to learning adaptations in preverbal 4-month-old infants. Concretely, monolingual infants display faster orientation towards their native language (Spanish or Catalan - Bosch & Sebastián-Gallés, 1997; French or American - Dehaene-Lambertz & Houston, 1998), while Spanish – Catalan bilingual infants show the opposite pattern, orienting faster to unfamiliar languages (Bosch & Sebastián-Gallés, 1997). Similarly, Spanish-Basque bilingual infants show longer sustained attention when processing their native languages and can distinguish them even if they belong to the same rhythmic class (Molnar et al., 2014). These results suggest that, from an early age, infants’ behavioural responses to familiar and unfamiliar languages are influenced by their linguistic context, with monolingual infants demonstrating a preference for their native language and bilingual infants displaying distinct attention patterns to familiar and unfamiliar language inputs, indicating early learning adaptations based on linguistic exposure. It has been proposed that bilingual language exposure might elicit cognitive adaptations that allow infants to accommodate the increased complexity of their linguistic environment. Distinctive attention allocation skills or an increased perceptual sensitivity are examples of the proposed adaptations (Byers-Heinlein & Fennell, 2014; Costa & Sebastián-Gallés, 2014; Kovács, 2015). In the current study, we test whether the previously observed behavioural adaptations are also reflected in distinct neural signatures for speech processing, specifically focusing on the language that is common across monolingual and bilingual infants.

Current neuroimaging evidence assessing the impact of early bilingual experience in the brain mechanisms underpinning speech processing is limited, especially in the age group of interest for this work (i.e., 4-month-olds), but results from these studies suggest that bilingual environments may impact early neural specialization. Nacar Garcia et al., (2018) demonstrated early neural specialization during language discrimination in 4.5-month-old monolingual infants (Spanish or Catalan), with these infants showing faster neural discrimination responses to their native language compared to an unfamiliar one from a different rhythmic class, which was in line with previous findings (Friederici et al., 2007). In contrast, bilingual infants (Spanish-Catalan) showed comparable responses for familiar and unfamiliar languages. Interestingly, brain responses to the shared native language (Spanish or Catalan) did not differ between monolingual and bilingual infants, suggesting that potential effects due to familiarity with the language may not have played a significant role. Another study used fNIRS to investigate the brain responses to spoken and signed language in English monolingual, unimodal bilingual and bimodal bilingual 4-to 8-month-old infants (Mercure et al., 2020). This study revealed no lateralization effects in monolingual and bimodal bilingual infants, whereas unimodal bilingual infants’ brain responses were right lateralized over posterior temporal regions for spoken and signed language conditions. The findings observed in Mercure et al., (2020) suggest that unimodal bilingual experience might influence early neural specialization, potentially due to the increased complexity of separating two spoken languages, as opposed to one spoken and one signed language. Differential developmental patterns in the neural responses to phonetic processing have been also identified across monolingual and bilingual infants (Petitto et al., 2012). Specifically, at 4 months of age, similar responses were observed between monolingual and bilingual infants in the inferior frontal region, which was the primary region showing sensitivity to the experimental manipulation involving native vs. non-native phonetic contrasts. However, at 12 months of age, bilingual infants exhibited perceptual sensitivity to both native and non-native phonetic contrasts in the left inferior frontal regions, in contrast to their monolingual counterparts who primarily showed sensitivity to native phonetic contrasts in this region (Petitto et al., 2012). In summary, neuroimaging evidence revealed specialized brain responses in monolingual infants for their native language and the recruitment of additional brain regions during speech processing in bilingual infants. In addition, these studies do not seem to provide evidence supporting an impact of language familiarity resulting from a reduced exposure to each of their native languages in bilingual infants. While some previous studies in monolingual newborns and young infants showed increased activation to familiar compared to unfamiliar languages (Minagawa-Kawai et al., 2011; Sato et al., 2012), the evidence for this effect appears inconsistent in infants aged 0-4 months of age, as other studies have not supported this outcome (Fava et al., 2014; May et al., 2011, 2018; Peña et al., 2010)

The aim of this study is to investigate whether 4-month-old monolingual vs. bilingual infants exhibit functional brain adaptations when processing speech on their shared native language. During the first year of life neuroplasticity is at its peak, making this period particularly relevant for studying the neural consequences of simultaneously acquiring two languages (Werker & Hensch, 2015). Language acquisition begins as soon as infants are able to hear spoken language, about 3 months prior to birth (e.g., Werker, 2018) and recent research has demonstrated the emergence of early brain specialization for prenatally heard language in monolingual newborns (Mariani et al., 2023). The earliest cognitive adaptations due to a bilingual environment have been observed in infants at 4 months of age (Bosch & Sebastián-Gallés, 1997; Molnar et al., 2014). However, it is unclear if the observed cognitive adaptations also reflect changes in the brain mechanisms underlying speech processing in bilingual infants at this young age, as this specific question has not been previously investigated. In a previous study we demonstrated a comparable functional brain network organization across bilingual and monolingual 4-month-olds from the same population during a task-free condition (Blanco et al., 2021, 2022). The extent to which neural adaptations can be observed in response to speech stimuli in these two populations remains uncertain. The present research aims to address this gap by investigating the impact of early bilingualism on the neural systems supporting speech processing in preverbal infants. Based on the reviewed evidence, we predict a more functionally specialized network for speech processing in monolingual infants, potentially involving a left hemispheric dominance. In bilingual infants, more widespread brain responses and an increased participation of right temporal regions is expected.

This study was conducted in the Gipuzkoa region of the Basque Country (Spain), where Spanish and Basque languages co-exist. Studying language acquisition in a context where monolingual and bilingual families are similarly prevalent provides a unique opportunity to study the effect of early language experience on the brain mechanisms involved on speech processing. We measured the brain responses to Spanish forward (FW) and backward (BW) speech stimuli in 4-month-old Spanish-Basque bilingual and Spanish monolingual infants using functional near-infrared spectroscopy (fNIRS). Brain regions involved in spoken language processing are similar across infants and adults, with left temporal and frontal perisylvian brain regions playing a central role in processing speech sounds (e.g., Friederici, 2002; Meyer et al., 2002). Specifically, the primary and secondary auditory cortices in the superior temporal gyrus and the inferior frontal gyrus have shown activation in studies assessing brain responses to speech in newborns (Perani et al., 2011) and older preverbal infants (Dehaene-Lambertz et al., 2002; Shultz et al., 2014). However, infants’ brain activation patterns for FW and BW speech stimuli described in the fNIRS literature show high variability in terms of the lateralization, shape, and direction of the observed hemodynamic responses (Grossmann et al., 2010; Kotilahti et al., 2010; May et al., 2011; Zhang et al., 2022, for a review see Issard & Gervain, 2018) and in the consistency of the observed responses across oxy- and deoxyhemoglobin (HbO and HbR) parameters (Mercure et al., 2020; Minagawa-Kawai et al., 2011; Peña et al., 2003). Here, to maximize data quality and reduce attrition rate, acquisition of fNIRS measurements were performed during natural sleep. We used a stimuli randomization approach optimized for general linear model (GLM) analysis that facilitates the acquisition of a large number of trials per participant while reducing testing time (Kao et al., 2009), which additionally improved our study’s statistical efficiency (Chen et al., 2022). This methodological approach allowed us to collect a sample of 57 infants, with a large number of trials per participant (>17 trials per condition) and optimal data quality, to accurately characterize 4-month-old monolingual and bilingual infants’ brain hemodynamic responses during speech processing.

## Methods

### Participants

Eighty-one 4-month-old infants participated in this study (Ethnicity: white). Participants were recruited from the same region of the Basque Country (Gipuzkoa), a predominantly bilingual region in which Spanish and Basque are learnt at home or/and at nursery school from a very young age. A socioeconomic status questionnaire was completed to ensure that families showed similar levels of education, parental occupation, and household income. Participants’ percentage of exposure to Spanish and Basque during the first months of life was assessed with a questionnaire filled by the parents. Two language groups were considered: Spanish–Basque bilingual (BIL) infants and Spanish dominant (SP) infants. Infants raised in a Spanish-Basque bilingual environment, those that were exposed to their two native languages from birth, with a percentage of exposure to Spanish between 20% and 80%, formed the bilingual group. Participants with an exposure to Spanish higher than 80% of their time were included in SP group (**Table 1**).

**Table 1.** Summary information of the participants included in the study.

| Language background | n | Mean age (days) | Exposure to Spanish (%)<br>mean ± SD | Range [max - min] (%) |
| --- | --- | --- | --- | --- |
| Spanish-Basque (BIL) | 26 | 125 | 53 ± 15.9 | [20.4 - 78.6] |
| Spanish (SP) | 31 | 124 | 94 ± 6.9 | [80.5 - 100] |

In nine of the 81 infants no recording took place as they were unable to fall sleep (n = 6 BIL, n = 3 SP). Fourteen infants (n = 7 BIL, n = 7 SP) were excluded for presenting insufficient data quality, as determined by the data quality assessment routine during preprocessing. One participant was excluded for presenting a regular exposure to Galician language. Fifty-seven participants were included in the final sample, for which data was analysed and results are presented: 26 BIL infants (15 girls; mean age = 125 days; exposure to Spanish *M* = 53%, *SD* = 16%, range = [20.4% - 78.6%]), 31 SP infants (16 girls; mean age = 124 days; exposure to Spanish *M* = 94%, *SD* = 6.9%, range = [80.5% - 100%]). This study was carried out at the Basque Center on Cognition, Brain and Language (BCBL) and received approval from its local ethical committee. The study involved the participation of infant subjects. Parents were informed about the aim of the study, the experimental procedures, and their right to withdraw from the study at any moment without providing a reason and with no negative consequences. Written informed consent was obtained from the parents prior to data acquisition.

### Stimuli

Three Spanish native female speakers were recorded reading aloud sentences from Saint-Exupéry’s *The Little Prince* book. Each speaker recorded the same 120 utterances which ranged in number of syllables between 15 and 18 (30 utterances of each length). From this set, 24 utterances were selected for each speaker, with no repetition between them. Sentences were manually segmented to the precise onset and offset time, and next normalized in intensity (70 dB) and peak amplitude across speakers using *Praat* (Boersma & Weenink, 2007). The final set comprised 72 utterances with a mean duration of 3.2 ± 0.2 seconds, range [3 – 3.7]. Backward (BW) sentences were generated from the set of forward (FW) sentences using *Praat*. The paradigm included 24 blocks per condition (i.e., FW and BW speech stimuli). Each block was formed by three sentences of the same speaker, with an inter-sentence interval of 0.5 seconds **(Fig.1a)**.

**Fig. 1.**
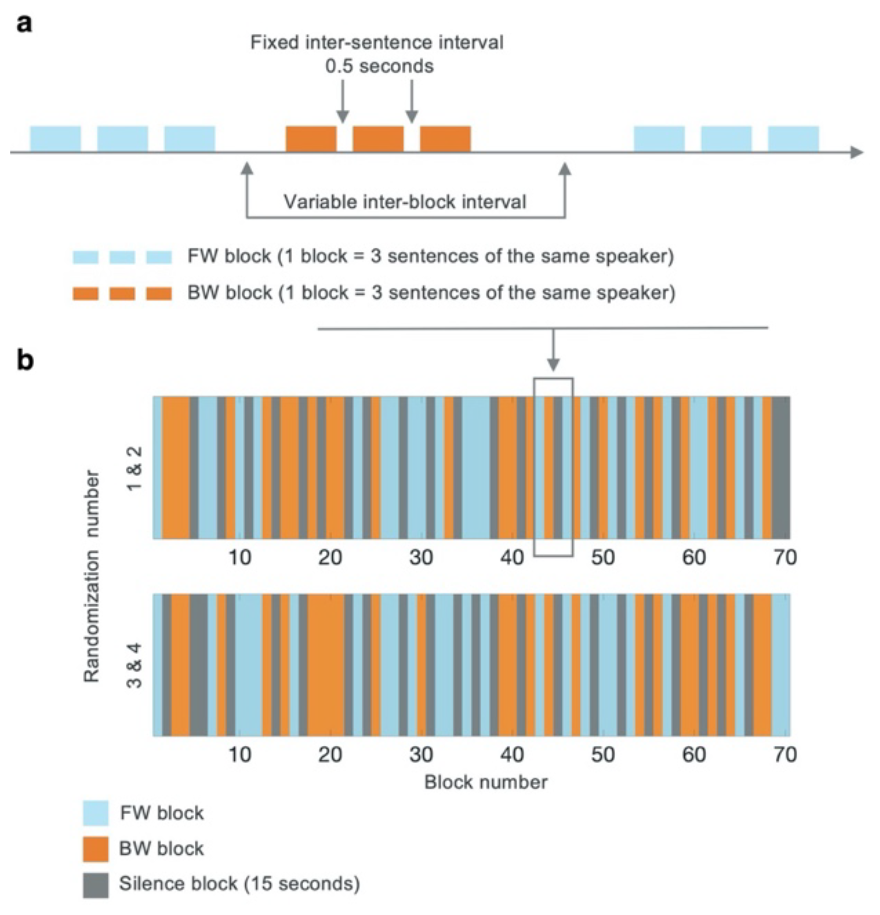
Illustration of the stimuli presentation procedure used in this study. **a)** Blocks for FW and BW conditions were formed by three individual sentences separated by an interval of 0.5 seconds. **b)** The experimental designs for the four stimuli randomization paradigms employed in the current study which were optimized for detection power and estimation efficiency (two additional paradigms were created by swapping the FW and BW conditions).

An experimental design optimized for the detection of brain activation and for the estimation of the shape of the hemodynamic response was implemented (Kao et al., 2009). Briefly, the main goals in task-based fNIRS studies measuring the brain responses to stimuli are two: 1) detecting brain regions that show activation elicited by the presentation of the stimuli (i.e., detection power), and 2) providing an accurate estimation of the shape of the hemodynamic response (i.e., estimation efficiency). However, pursuing both goals simultaneously in one study is also possible with an optimal multi-objective randomized experimental design (Kao et al., 2009). This method requires the user to specify a weight for four design objectives: i) detection power, ii) estimation efficiency, iii) desired fraction of trials per condition (i.e., stimulus frequency), and iv) predictability (i.e., controlling psychological confounds induced by subject’s prediction of the successive events). Based on the goals of the experiment, the algorithm searches an optimal design sequence via a multi-objective optimization problem that is solved using a genetic algorithm. The initial search pool is formed by a set of experimental designs that are known to be optimal for each of the objectives independently, and a set of random experimental designs. Then, the algorithm proceeds iteratively generating new design sequences using three different methods [*crossover* – interchange portions of different sequences; *mutation* – randomly replace specific events (i.e., stimulus, rest); *immigration* – add new designs sequences] and assessing the fitness of each design for each experimental objective. In each iteration the most optimal designs are stored, as they are used to create the initial pool of the subsequent iteration until a stopping rule is met (e.g., desired multi-objective design efficiency achieved). Two design sequences were generated using this approach **(Fig. 1b)**, and two additional design sequences were generated by swapping the order of the experimental conditions (i.e., FW and BW speech stimuli) on the first two design sequences. Participants were randomly assigned to one of these four randomizations, all of them with the same number of blocks per condition (i.e., 24 blocks) and a similar duration of approximately 17 minutes.

### Data acquisition procedure

FNIRS measurements were performed with a NIRScout system (NIRx Medical Technologies, CA, USA) at wavelengths 760 and 850 nm with a sampling frequency of 15.625 Hz. Eight light emitters and 12 detectors were positioned on a stretchy fabric cap (Easycap GmbH, Germany) over frontal, temporal, and parietal regions of both hemispheres according to the international 10-20 system. Nasion, inion and preauricular points were used as external head landmarks. Each pair of an adjacent light emitter and a detector formed a single measurement channel, which generated 24 channels for each haemoglobin oxygenation state (i.e., oxyhaemoglobin, HbO and deoxyhaemoglobin, HbR) consisting of source-detector separation distances ranging from 20 to 30 mm **(Fig. 2)**.

**Fig. 2.**
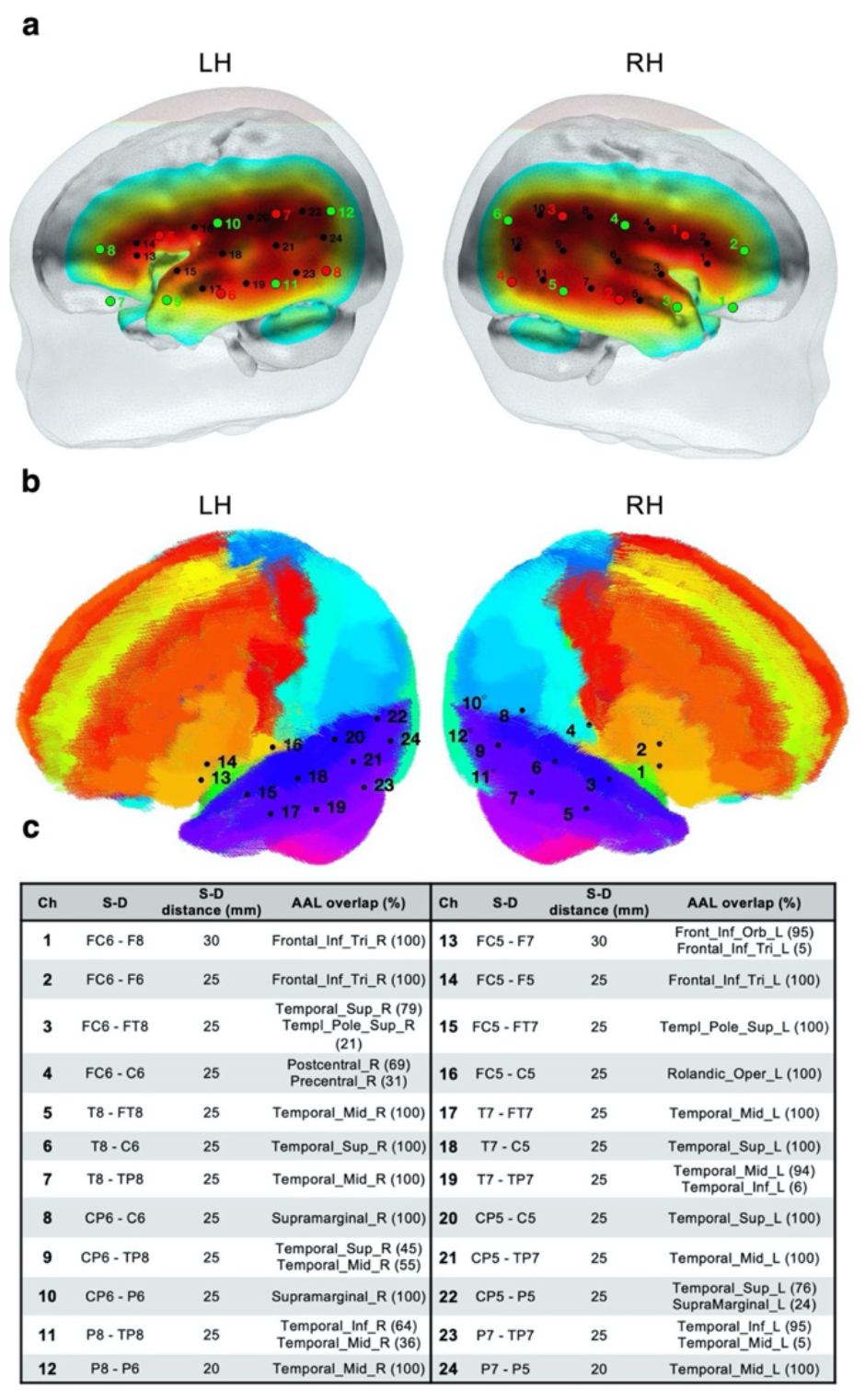
**a)** fNIRS optode (sources in red, detectors in green) and channel (black) localization in the current experimental setup. The normalized sensitivity profile of the current optode configuration is displayed in a 6-month-old infant head model. **b)** fNIRS channel localization registered to a 6-month-old infant AAL template. **c)** Brain labels of the fNIRS channels based on the probabilistic spatial registration of the fNIRS optode setup to a 6-month-old infant AAL template. Ch = Channel; S = Source; D = Detector.

The sensitivity profile of the fNIRS probe setup was computed to provide information of the brain areas under investigation, and for the result visualization purposes. The probe setup (i.e., sources and detectors) was registered to an average 6-month-old infant template (Richards et al., 2016) to compute the sensitivity matrix of the source-detector configuration using *Toast++* (Schweiger & Arridge, 2014). The aggregated sensitivity profile of the fNIRS probe was obtained by adding the normalized cortical sensitivity profiles of each individual channel **(Fig. 2)**. Channel positions were defined as the grey matter node which coordinates were closest to the central point of the maximum sensitivity path along each source-detector pair. A 6-month-old average atlas (Akiyama et al., 2013) was used to compute a probabilistic spatial registration of the cortical structures underlying each channel **(Fig. 2)**. Channel coordinates were first transformed from the average 6-month-old template employed for computing the sensitivity profile to the Akiyama et al., (2013) average T1 template space using Advanced Normalization Tools (ANTs, Avants et al., 2009). Then, coordinates were registered to the Akiyama et al., (2013) anatomic atlas, formed by 116 cortical regions based on Automated Anatomical Labeling (AAL, Tzourio-Mazoyer et al., 2002). For each channel, the AAL anatomical labels within 20 mm were defined, and the percentage of overlap with each AAL region was calculated **(Fig. 2)**.

During the study infants rested on their parents’ lap facing the loudspeakers presenting the stimuli. First, the fNIRS cap was placed on the infants’ head. Then, a feed and wrap approach was used to promote sleep. The experiment began when clear signs of sleep were noticeable on the infant (e.g., eyes closed, lack of movements). During the pilot phase of this study, it was observed that the sudden change from a room in complete silence to high-volume stimuli presentation was making the infants awake. Therefore, for the next sessions it was decided to progressively increase volume during the first 30 seconds manually by one of the experimenters to avoid this effect. This portion of the data was excluded from the analyses during data preprocessing.

### Data preprocessing

All data preprocessing and analyses were computed in MATLAB (R2012b, R2014b, Mathworks, Massachusetts) using in-house scripts as well as third-party toolboxes and functions. First, light intensity data (i.e., raw data measured at the instrument) were converted into optical density changes by computing the negative logarithm of the ratio between detected light intensity at each time point and a reference baseline value (i.e., the mean signal). The first 30 seconds of the experimental task were censored as during this period the volume progressively increased towards the maximum volume level. Motion artifacts were identified using an automated function from Homer2 fNIRS processing package (Huppert et al., 2009). Time points identified as motion on this step were censored and excluded from further analysis. Other motion induced spikes and signal drifts were corrected using a wavelet-based despiking method adapted for fNIRS (Patel et al., 2014). Optical density data were converted into HbO and HbR concentration changes by means of the modified Beer-Lambert law with differential path length factors of 5.3 and 4.2 (Scholkmann & Wolf, 2013). The contribution of high-frequency physiological noise sources (e.g., respiration and cardiac pulse) was attenuated by means of a zero-phase low-pass filter with cut-off frequency 0.5 Hz. Slow frequency fluctuations and signal drifts were modelled by up to 8^th^ order Legendre polynomials. Globally occurring hemodynamic processes in cerebral and extracerebral tissues assumed to largely reflect systemic hemodynamic changes were also included in the nuisance regression model. Prior to this step the average signal was low pass filtered with the same parameters as the data to avoid reintroducing frequencies of non-interest. As HbO and HbR are differently affected by global systemic processes, data of each haemoglobin chromophore were filtered independently by including in the model either the global HbO or HbR signal. Additional information about the parameters used in the preprocessing pipeline and about the effect of applying global signal regression are available in **supplementary materials**.

### Data analysis

For each participant, and for each experimental condition, the percentage of block-related data included for data analysis after censoring motion affected time points was calculated. This allowed us to assess that every time point inside each block (i.e., pre-stimulus baseline and stimulus block duration, around sixteen seconds in total) retained a similar number of data points after data censoring. The aim was to ensure that, for both experimental conditions, time points during the entire block were similarly represented to accurately model the hemodynamic response. In the BIL group, the mean percentage of block-related time completed for the FW condition was 94 % ± 4 %, range = [82 % - 99 %], and for the BW condition this percentage was 94 % ± 3 %, range = [88 % - 99 %]. In the SP group, the mean percentage of block-related data included for the FW condition was 92 % ± 6 %, range = [72 % - 99 %], and for the BW condition this percentage was 92 % ± 5 %, range = [80 % - 99 %]. These results imply that at least 17 out of 24 blocks (72%) were completed by each infant and for each condition, although in most infants the number of completed blocks was indeed higher. This analysis also confirmed that all time points inside each block, and for FW and BW conditions, were similarly represented and no data segment was particularly affected by the censoring procedure **(supplementary materials)**.

Two general linear model (GLM) based analyses were performed to study the brain’s responses to FW and BW speech stimuli. The first GLM analysis aimed to detect areas/channels showing functional activation to the presentation of FW and BW speech stimuli. Here the operational definition of functional activation for fNIRS based studies (i.e., increase in HbO and a decrease in HbR) is considered (Obrig & Villringer, 2003). Regressors were generated for each experimental condition by convolving boxcar functions of average trial duration (i.e., eleven seconds) with a model of the hemodynamic response function, a gamma function with peak at four seconds **(supplementary materials)**. The peak of the gamma function was selected based on comparisons with the hemodynamic response estimated in the second GLM analysis implemented in this study **(supplementary materials)**, which is described below. In addition, boxcar regressors were included to model potential baseline differences between each data segment formed after removing censored time periods. This analysis yielded the *β*-values representing the mean effect of individual functional responses to each experimental condition (i.e., FW and BW speech) for HbO and HbR.

Statistical analyses were performed on the estimated *β*-values. First, one-sample t-tests were performed on individual *β*-values to detect brain regions (i.e., channels) that were sensitive to each experimental condition within each group (i.e., BIL, SP). For this analysis, statistical tests were corrected for multiple comparisons using the false discovery rate (FDR) method (q < 0.05, Benjamini & Hochberg, 1995). Second, for the statistical comparisons between groups, differences in mean *β*-values between BIL and SP infants were modelled in a two-way mixed effects ANOVA for each channel, for HbO and HbR separately, and corrected for multiple comparisons using the false discovery rate (FDR) method (q < 0.05, Benjamini & Hochberg, 1995). The mixed design was determined by the two independent variables that were manipulated in this experiment, one repeated-measures independent variable with two levels (i.e., FW and BW conditions), and one between-group independent variable with two levels (i.e., language background BIL and SP). Post-hoc t-tests corrected for multiple comparisons were conducted using Tukey’s method.

The second GLM analysis aimed at estimating the shape of the hemodynamic response evoked by each stimulus condition, which can be accomplished by using a finite impulse response or deconvolution model. For each infant, HbO and HbR channel time courses were modelled using a Gaussian basis set consisting of one-second gaussian functions, spanning [-5 to 25] seconds around stimulus onset. Potential baseline differences between each of the individual periods generated by motion censoring were modelled using boxcar regressors. Hemodynamic responses for FW and BW conditions were described on each group to assess their consistency with the expected characteristics of the hemodynamic response extracted from fNIRS signals (Obrig & Villringer, 2003), but these curves were not used for statistical comparisons.

## Results

### Group-level hemodynamic responses in bilingual and Spanish infants

On each group (i.e., BIL, SP) significant responses were determined by means of one-sample t-tests computed on the *β*-values on each channel and for each condition (FW, BW), which were corrected for multiple comparisons using the FDR method (q < 0.05, Benjamini & Hochberg, 1995). We highlight results that survived multiple comparisons correction, but present also uncorrected results for completeness (Taylor et al., 2023). Complete channel statistics for BIL and SP groups, and for FW and BW conditions are reported in **supplementary materials**. For the FW condition, BIL infants displayed significant brain responses in channels located in bilateral superior and middle temporal regions, and in bilateral inferior frontal regions (**Fig. 3a**). Activation responses were observed in bilateral middle and superior temporal regions. In contrast, channels located in bilateral inferior frontal regions showed significant deactivation responses consisting of hemodynamic responses following an inverted pattern (i.e., a decrease in the local concentration of HbO and an increase in HbR). SP infants also showed activation responses on channels located in left superior and middle temporal regions and in one channel located in right middle temporal region (**Fig. 4a**). SP infants displayed deactivation responses right inferior frontal regions. In both groups, channels exhibiting significant responses were consistent with the estimated hemodynamic responses obtained with the deconvolution model (**Fig. 3b and Fig. 4b**). Moreover, the estimated group-level hemodynamic responses matched the operational definition of functional activation for fNIRS studies (i.e., an increase in the local concentration of HbO and a decrease in HbR, or the opposite), supporting the reliability of the obtained outcomes.

**Fig. 3.**
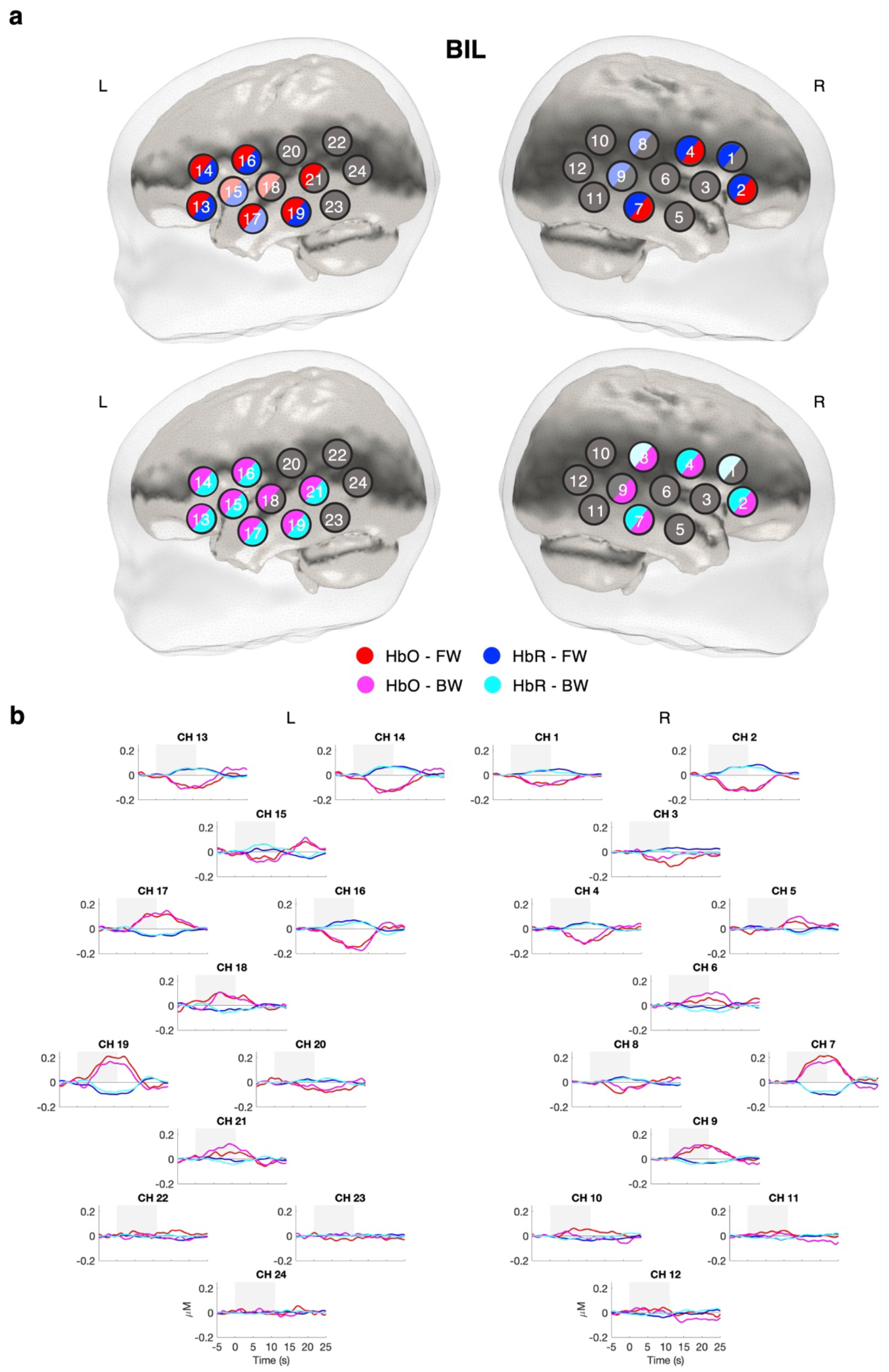
a) Channels showing a significant activation/deactivation response as determined by one-sample t-tests for FW (top) and BW (bottom) conditions in BIL infants (FDR corrected, q<0.05). Uncorrected results (p<0.05) are displayed in a lighter colour. b) Estimated group-level hemodynamic responses at each channel location and for each experimental condition. Time zero indicates stimulus onset.

**Fig. 4.**
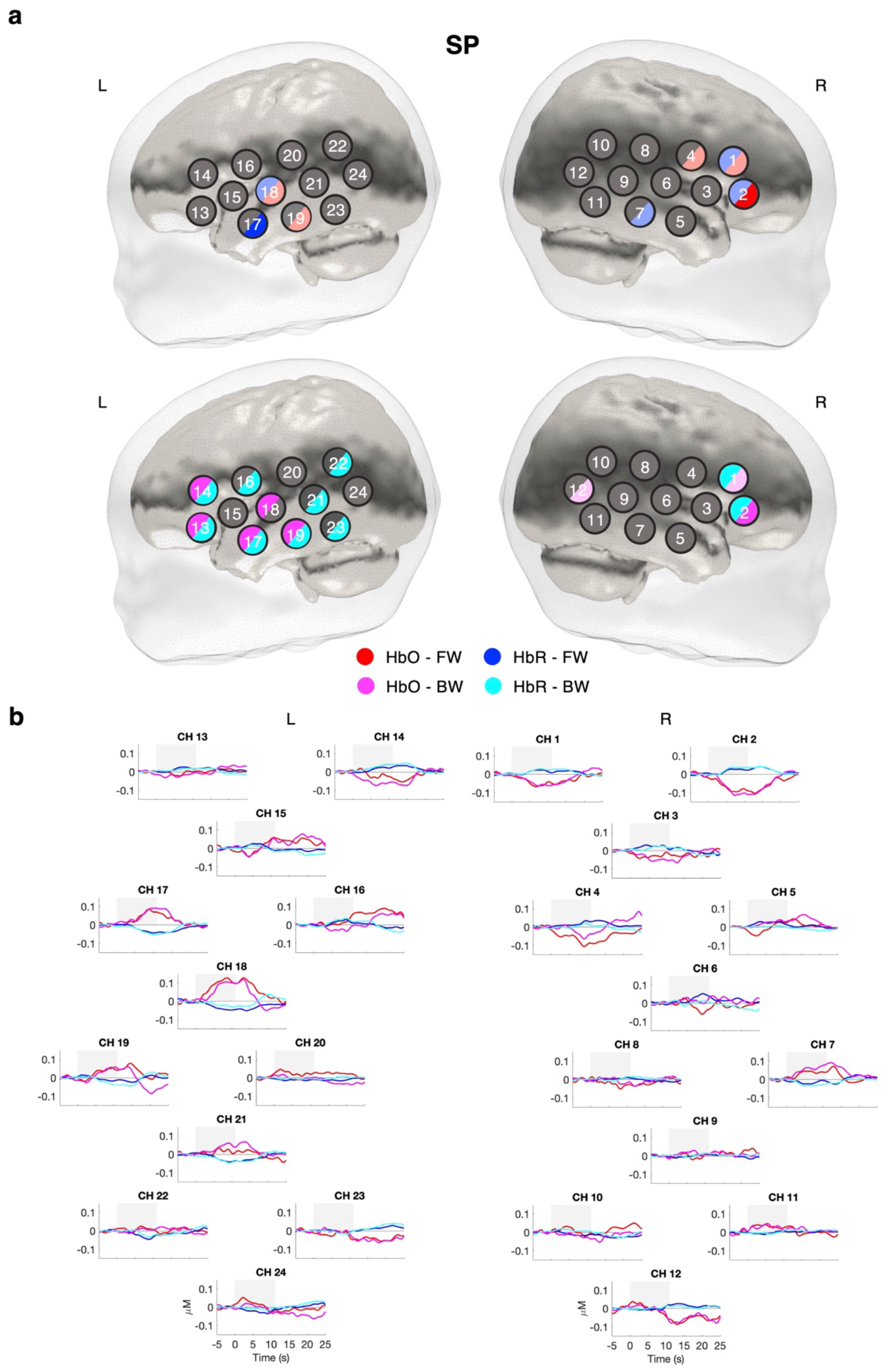
a) Channels showing a significant activation/deactivation response as determined by one-sample t-tests for FW (top) and BW (bottom) conditions in SP infants (FDR corrected, q<0.05). Uncorrected results (p<0.05) are displayed in a lighter colour. b) Estimated group-level hemodynamic responses at each channel location and for each experimental condition. Time zero indicates stimulus onset.

### Comparisons between bilingual and Spanish infants

Significant results obtained from the two-way mixed ANOVA (corrected and uncorrected) are reported in the main text (**Table 2**). Complete statistical results are included in **supplementary materials**. A main effect of Condition (FW, BW) was observed in one channel located in the left inferior frontal region. In this channel the BW condition displayed a stronger response as compared to the FW condition. A main effect of Language (BIL, SP) revealed stronger responses in BIL as compared to SP infants in channels located in the left inferior frontal region and in bilateral middle temporal regions (**Table 2**). Concretely, BIL infants displayed stronger activation responses in bilateral middle and superior temporal channels. BIL also showed significantly stronger deactivation responses than SP infants in left inferior frontal channels. Interaction contrasts showed significant effects in two channels (**Table 2, Fig. 5a**). In channel 16, located in the left inferior frontal region, there was a significant interaction between infants’ language background and experimental condition (HbR, F_1, 55_ = 11.899, p = 0.001). Post-hoc tests in channel 16 indicated stronger deactivation responses in BIL infants than in SP infants for the FW condition (HbR, t = 3.528, p = 0. 004, d = 0.938, 95% CI [0.030, 0.225], **Fig. 5b** and **Fig. 5c**) and stronger deactivation responses in SP infants for the BW as compared to the FW condition (HbR, t = −2.981, p = 0.022, d = 0.505, 95% CI [-0.131, −0.006]). In channel 19, located in the left middle temporal region, a significant interaction between language background and experimental condition was observed (HbO, F_1, 55_ = 4.28, p = 0.043; HbR, F_1, 55_ = 10.55, p = 0.002). In channel 19 group differences were characterized by stronger activation responses for FW speech in BIL infants as compared to SP infants (HbO, t = 3.345, p = 0.007, d = 0.890, 95% CI [0.069, 0.659]; HbR, t = −4.073, p < 0.001, d = 1.083, 95% CI [-0.328, −0.066]) and by stronger activation responses in SP infants for the BW as compared to the FW condition (HbR, t = 2.833, p = 0.026, d = 0.353, 95% CI [0.002, 0.126]).

**Table 2.**
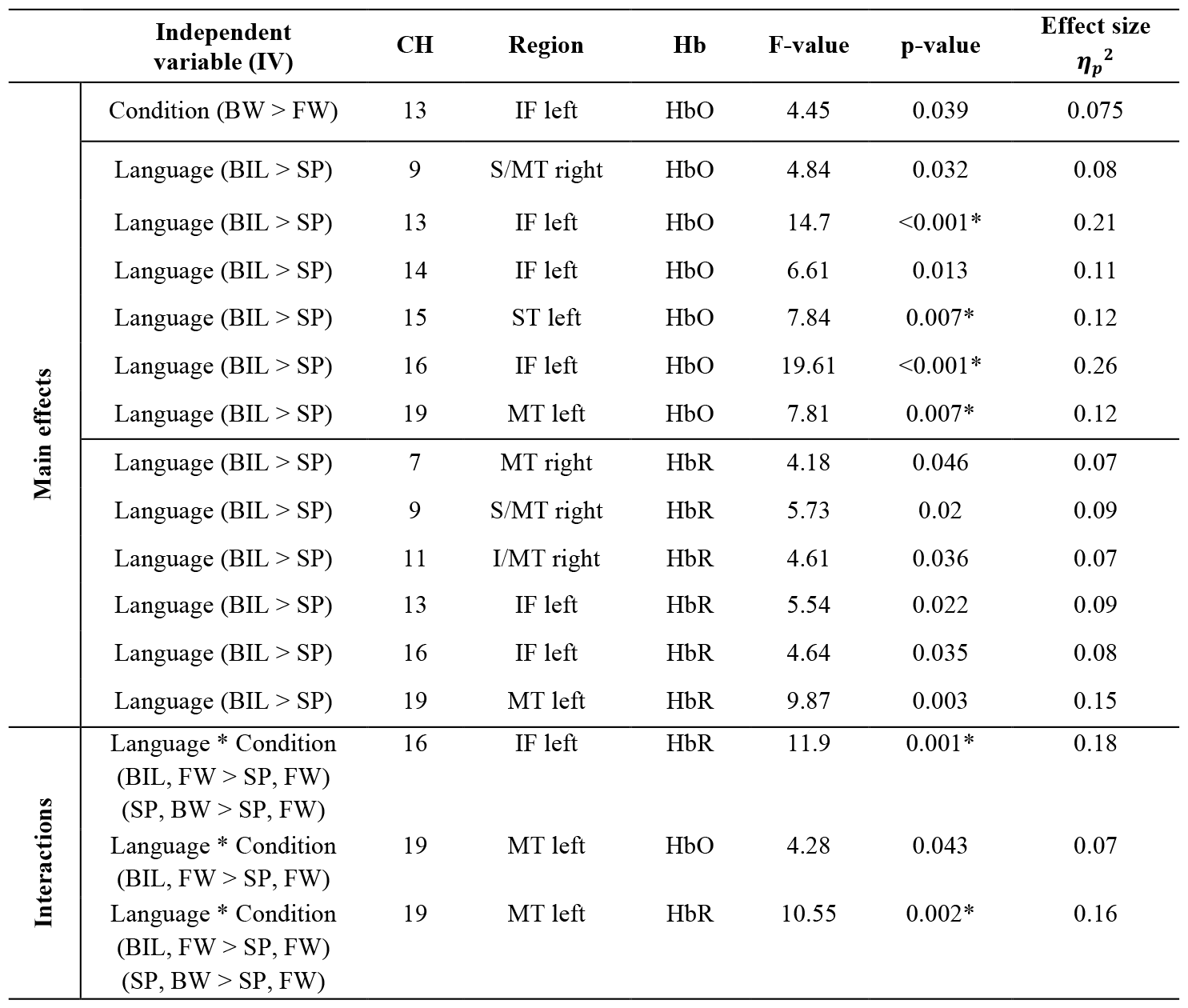
Results from the two-way mixed ANOVA including main effects and interactions where significant differences between experimental groups were observed. ^*^ Denotes effects that survived multiple comparisons correction (FDR method). Language (BIL, SP), condition (FW, BW). S = superior, M = middle, T = temporal, F = frontal, I = inferior.

**Fig. 5.**
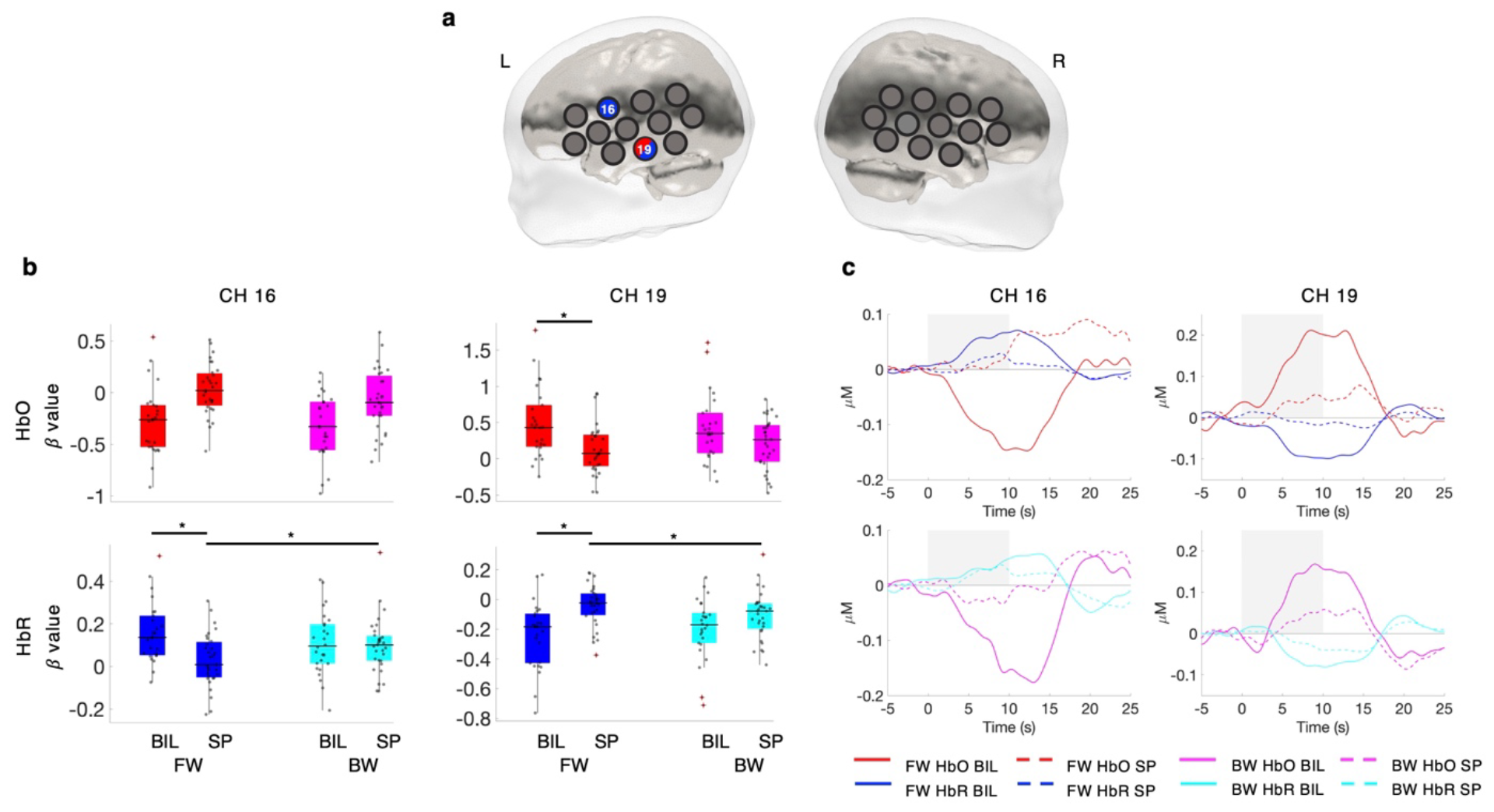
a) Channels 16 (HbR) and 19 (HbO and HbR) showed a significant interaction effect between infants’ language background (BIL, SP) and experimental condition (FW, BW). b) Boxplots of post-hoc tests computed in channels 16 and 19. ^*^ Denotes significant effects after multiple comparisons correction. c) Hemodynamic responses in BIL and SP infants for the FW and BW conditions in channels 16 and 19.

We also tested the effect of familiarity with Spanish language by including percentage of exposure to Spanish as a covariate in our model. For all channels and conditions (HbO and HbR), the effect of percentage of Spanish as covariate was only significant on channel 16 (HbR, F_1, 55_ = 4.564, p = 0.037), and including this covariate in the model did not substantially alter the direction of the observed effects on this channel without including the covariate [main effect of Language (BIL > SP), F_1, 55_ = 8.742, p = 0.005; interaction Language^*^Condition, F_1, 55_ = 4.22, p = 0.045].

## Discussion

This study used fNIRS to describe 4-month-old infants’ brain responses during speech processing and to compare these responses across infants growing up in a bilingual (Spanish and Basque) vs. a monolingual (Spanish) environment. The current results provide the first evidence of experience-dependent adaptations on the brain mechanisms underlying speech processing in bilingual infants as early as 4 months of age.

Within-group results showed that BIL and SP dominant infants displayed functional brain responses organized around perysilvian areas that are consistent with previous infant literature assessing speech processing (Dehaene-Lambertz et al., 2002, 2006). However, BIL and SP infants’ brain responses to speech stimuli exhibited unique characteristics. For both speech conditions (i.e., FW and BW speech), BIL infants showed a more widespread pattern of activation in bilateral middle/superior temporal and inferior frontal regions, and a larger number of channels showing significant responses to the presented speech stimuli as compared to SP infants. Furthermore, BIL infants displayed significant activation in channels located on the right middle and superior temporal regions, whereas SP infants did not exhibit significant activation in right temporal areas.

Direct statistical comparisons between BIL and SP infants’ brain responses demonstrated differences between these groups in various fNIRS channels. Bilingual infants demonstrated a generalized pattern of increased brain responses to both FW and BW speech as compared to SP infants, predominantly observed over left inferior frontal, and middle and superior temporal regions. An interaction effect in channels 16 and 19, localized in the left inferior frontal and middle temporal region respectively, revealed that BIL and SP infants were differentially sensitive to FW and BW speech stimuli in these channels. Both channels followed similar response patterns, with BIL infants showing larger responses to FW speech than SP infants, and SP infants showing enhanced responses to BW as compared to FW speech. In summary, more widespread responses, increased participation of right temporal areas and similar response patterns across FW and BW speech were evident in BIL infants.

Spanish dominant infants showed brain responses in temporal regions that were mostly confined to the left hemisphere, and they displayed significantly stronger responses for BW as compared to FW speech in channels localized in left inferior frontal and left middle temporal regions (channels 16 and 19). Previous evidence assessing left hemispheric superiority in monolingual infants for speech processing is mixed. While some studies report left lateralized responses to native speech stimuli in neonates (Peña et al., 2003; Sato et al., 2012) and 3–4-month-old infants (Dehaene-Lambertz et al., 2002; Minagawa-Kawai et al., 2011), other works observed bilateral response patterns in neonates (May et al., 2011, 2018; Perani et al., 2011) and older, 3-to 6-month-old, infants (Homae et al., 2006; Mercure et al., 2020; Zhang et al., 2022). Moreover, our findings demonstrate that 4-month-old SP infants are sufficiently specialized in certain characteristics of their native language to show differential brain responses when compared to other speech-like stimuli with similar features. In SP infants, areas within the left inferior frontal gyrus and the left middle temporal region were distinctively sensitive to the properties of FW and BW speech. Neuroimaging studies on preverbal infants have shown specialized brain responses to speech conditions such as BW or flattened speech (Dehaene-Lambertz et al., 2002; Homae et al., 2006; May et al., 2018; Perani et al., 2011). Note that other works with 5-6 month-old monolingual infants have reported no differences in the brain responses to FW vs. BW speech (Zhang et al., 2022) or to familiar vs. unfamiliar languages (Mercure et al., 2020). Typically, studies have reported greater activation for FW compared to BW speech, which might indicate preference due to familiarity, especially in newborns (Mehler et al., 1988). In contrast, our results indicate a stronger response to BW as compared to FW speech in SP infants.

Infants raised in a bilingual environment display a distinct and broader pattern of brain activity in comparison to their monolingual peers. Specifically, bilingual infants show enhanced activity in the left inferior frontal and bilateral middle and superior temporal regions. Prior research has indicated differing brain activity patterns in older bilingual infants, particularly an increased engagement of the right hemisphere during linguistic tasks (Ferjan Ramírez et al., 2017; Mercure et al., 2020). This expansive engagement of the speech processing network in bilinguals, especially the pronounced activity in the right hemisphere’s temporal regions, mirrors observations from studies on bilingual adults (Hull & Vaid, 2007; Pierce et al., 2015; Połczyńska et al., 2017; Sulpizio et al., 2020) and children (Jasinska & Petitto, 2013). Our findings show that bilingual experience modulates brain activity in key language regions, including the left inferior frontal, left middle, and superior temporal areas, and their counterparts in the right hemisphere, as early as 4 months old. The early emergence of bilingualism-induced adaptations suggests that even a brief four-month exposure to two languages can profoundly influence the neural mechanisms responsible for language processing

Our results also indicate that bilingual infants show similar patterns of brain activity for FW and BW speech. One possibility is that BIL infants show a protracted functional specialization to their native language due to the increased variability of a bilingual environment, and the reduced exposure to each of their languages (Costa & Sebastián-Gallés, 2014; Singh, 2021). A delayed functional specialization in bilingual infants might be indicative of a different developmental transition in the perceptual sensitive periods during speech processing (Kovács, 2015; Werker & Hensch, 2015). This adaptation might be advantageous in a context of reduced exposure to each language input, as it might help bilingual infants to maintain the perceptual sensitivity to certain linguistic contrasts when monolingual infants are no longer able to do so (Petitto et al., 2012), and achieve an effective neural discrimination for two languages as opposed to one at a later age (Garcia-Sierra et al., 2011). As far as authors are aware, no previous studies in the literature have investigated if bilingual infants are capable of distinguishing FW and BW speech in their native languages, nor bilingual infants’ brain responses have been compared across these conditions.

Bilingual infants at this age are able to behaviourally discriminate their two native languages, and their native language from an unfamiliar one. Specifically, at this age Spanish-Basque bilingual infants are able to discriminate their two native languages even though they belong to the same rhythmic class (Molnar et al., 2014). Similarly, Spanish-Catalan bilingual infants are able to distinguish a familiar (i.e., SP or Catalan) from an unfamiliar language from the same rhythmic class such as Italian (Bosch & Sebastián-Gallés, 1997). Here, we used BW speech as a non-linguistic acoustic control for FW speech. This condition preserves fast transitions present in FW speech and it also shows similar pitch and intensity characteristics. However, several segmental and suprasegmental information is altered in BW sentences, and this condition is distinctly non-linguistic since most of its sounds cannot be produced by the human vocal tract. Considering that in the first months of life infants rely mostly on prosodic cues for language discrimination (Mehler et al., 1988), it seems unlikely that BIL infants are not able to perform the distinction between FW and BW Spanish at this age.

A complementary explanation is that the observed findings might be related with adaptations in other domain-general processes such as attention and learning strategies in bilingual infants as an effect of an increased environmental complexity. Flexibility of attention and increased openness to novel information have been proposed as specific cognitive adaptations that might allow bilingual infants to cope with the increased complexity of a dual language environment (Bialystok, 2015; Costa & Sebastián-Gallés, 2014). Research in support of this view shows different allocation of attention to familiar and unfamiliar languages during language discrimination tasks (Bosch & Sebastián-Gallés, 1997; Molnar et al., 2014) and a differential attention to audio-visual cues (i.e., eyes and mouth) during language processing (Ayneto & Sebastian-Galles, 2017; Pons et al., 2015). Various works have also reported differences in attentional orienting processes between monolingual and bilingual infants, with bilingual infants showing a marked preference towards novel information (Arredondo et al., 2022; D’Souza et al., 2020; Kalashnikova et al., 2021; Sebastián-Gallés et al., 2012; Singh et al., 2015). Our outcomes show enhanced responses to BW speech in SP infants as compared to FW speech, while BIL infants show comparable responses across FW and BW conditions. This might suggest that SP infants demonstrate increased interest towards a novel/unfamiliar condition such as BW speech, while for BIL infants both inputs are perceived as equally salient. Adaptations that allow bilingual infants to attend novel information more readily might facilitate the acquisition of two languages as opposed to one in a context of increased linguistic variability.

We observed higher participation of left frontal and middle temporal regions in BIL infants during FW and BW speech conditions, and larger responses in these regions during BW speech in SP infants. It is challenging to disentangle the exact role played by these regions during our speech processing task. Both regions are part of the fronto-temporal network supporting language processing (Friederici, 2002), but they are also recruited during other domain-general cognitive functions outside the language domain which have been shown to be modulated by early bilingual experience (Arredondo et al., 2017, 2022). Follow-up studies with increased sample size should aim to investigate the interactions between these brain regions during speech processing tasks, for example by means of effective connectivity approaches, such as dynamic causal modelling (Friston et al., 2013; Tak et al., 2015), that enable investigating direct causal interactions between brain regions.

We accurately described brain responses to speech in 4-month-old infants using fNIRS. Consistent responses were observed across HbO and HbR, with hemodynamic responses following the canonical shape (Obrig & Villringer, 2003; Yücel et al., 2021). This indicates that potential confounding effects due to systemic physiology and/or motion artifacts were accurately reduced by our experimental design and signal processing pipeline, which included global signal regression to mitigate the effect of physiological confounds. In contrast to temporal regions, where canonical activation responses were observed, inferior frontal regions showed inverted or deactivation responses (i.e., decrease in HbO and increase in HbR). This effect was not induced by the global signal regression step applied during preprocessing, as hemodynamic responses computed independently without applying this step also showed the same pattern (**supplementary materials**). The effect was present across experimental groups and conditions, indicating that the underlying process was not impacting differently any of the groups, nor was it related with the presented speech stimuli. Furthermore, deactivation responses cannot be attributed to a generalized altered response pattern during sleep, as canonical hemodynamic responses were observed in bilateral temporal regions during the presentation of FW and BW speech stimuli. Deactivation responses have been reported in previous studies measuring brain responses to auditory stimuli in sleeping infants (Abboub et al., 2016; Cabrera & Gervain, 2020; Dehaene-Lambertz et al., 2006). Tentative explanations for this effect include habituation of brain responses due to stimulus repetition (Cabrera & Gervain, 2020), a response inhibition / sleep protection mechanism (Wilf et al., 2016; Zou et al., 2020), or a compensatory mechanism that decreases blood flow to adjacent regions of brain areas being activated (Shmuel et al., 2001). Studies specifically targeted to investigate this effect are essential to improve our understanding of the neurophysiological mechanisms underlying this phenomenon, and to provide a more complete and accurate interpretation of infant hemodynamic responses measured with fNIRS.

One of the main limitations of this study lies in the fact that it involved the presentation of stimuli on a single language (i.e., Spanish), the one that was familiar to both monolingual and bilingual infants. As a result, this study does not provide a comprehensive evaluation of the distinctiveness of brain responses between these two populations and the potential role of linguistic familiarity. In other words, the current design does not allow to disentangle if the differential brain responses during speech processing observed in bilingual infants are a consequence of bilingualism per se, or instead they arise as a result of varying degrees of familiarity with the tested language. The inclusion of a second language unfamiliar to both groups was originally considered, but it was deemed impractical for infant participants due to constraints related to the duration of the experimental paradigm. We also explored the inclusion of percentage of exposure to Spanish as a covariate in our analysis, but this only yielded limited effects in a single channel. In this age group, the impact of language familiarity appears to vary, as certain studies on monolingual infants have noted this effect (Minagawa-Kawai et al., 2011; Sato et al., 2012), while others have not (Fava et al., 2014; May et al., 2011, 2018; Peña et al., 2010). Additionally, studies with bilingual infants of a comparable age have failed to observe any discernible effect of language familiarity (Mercure et al., 2020; Nacar Garcia et al., 2018). Nonetheless, we acknowledge that the presentation of both familiar and unfamiliar languages to monolingual and bilingual infants would have provided a more complete investigation of infants’ brain responses during speech processing and highlight the need to address this question in future studies.

The recruitment phase for this study was affected by the COVID-19 pandemic and as a consequence the sample size was smaller than originally planned. Despite sample size being relatively large for an infant neuroimaging study, we acknowledge that it might still have impacted statistical power and increased response variability. Potentially, some of the described effects might have been more accurately and reliably identified if each experimental group would have had a larger number of participants. However, recent works have demonstrated the pivotal role of trial sample size for task-based neuroimaging studies, with trial sample size having nearly the same impact as participant sample size for a study’s statistical efficiency (Chen et al., 2022). Here, each infant was presented with 24 trials per condition, with all infants contributing data for at least 17 trials per condition, a number that largely exceeds previous infant fNIRS studies and which might have assisted to partially alleviate this issue.

Infants’ sleep state might have also played a role on the differences in brain activation observed between monolingual and bilingual infants. All infants were scanned shortly after falling sleep, and thus it was assumed that all of them were in the same sleep state during fNIRS data collection. However, it is recognized that changes in autonomic physiology and arousal occur during sleep (Duyn et al., 2020), as well as differences in the brain responses to auditory stimuli (Dehaene-Lambertz et al., 2002; Kotilahti et al., 2005; Taga et al., 2018) and functional connectivity (Lee et al., 2020; Uchitel et al., 2023). Despite its inherent challenges, introducing additional neuromonitoring equipment for assessing sleep state in this population, for example electroencephalography or behavioural measures such as video recordings, would help future research to provide a more comprehensive understanding of infants’ functional brain responses during natural sleep.

## Conclusion

The current study offers compelling evidence of bilingualism-induced brain adaptations in 4-month-old infants during speech processing, supporting the notion that the neural foundations of bilingual language acquisition are established very early in life. Bilingual infants were found to engage bilateral inferior frontal and temporal regions during speech processing, exhibiting similar brain responses across speech modalities (i.e., FW and BW speech). In contrast, monolingual infants predominantly engaged areas in the left hemisphere and displayed specialized brain responses to each speech condition. These findings highlight the pivotal role of early language experience in shaping brain plasticity during early development. Specifically, an environmental factor such as bilingualism appears to trigger functional brain adaptations in early infancy, which might reflect the necessary learning adaptations that need to take place in bilingual infants to facilitate language acquisition in a more complex linguistic environment.

## Competing interest information

The authors declare no competing interest.

## Funding information

**B.B:** Basque Government Predoctoral Fellowship (2016-2020, PRE_2018_2_0154)

**M.M:** Spanish Ministry of Economy and Competitiveness (PSI2014-5452-P) and Natural Sciences and Engineering Research Council of Canada (Nos. 506948 and 506993)

**I.A:** Basque Government Predoctoral Fellowship (2020-2024, PRE_2022_2_0139)

**C.CG:** Spanish Ministry of Economy and Competitiveness (Ramon y Cajal Fellowship, RYC-2017–21845), the Spanish State Research Agency (BCBL “Severo Ochoa” excellence accreditation CEX2020-001010/AEI/10.13039/501100011033) and the Basque Government (BERC 2022–2025).

**M.C:** The Spanish State Research Agency (BCBL “Severo Ochoa” excellence accreditation CEX2020-001010/AEI/10.13039/501100011033) and the Basque Government (BERC 2022–2025).

## Acknowledgments

The authors would like to thank all the parents and infants who generously participate in our studies. The authors also would like to thank Elena Aguirrebengoa for her assistance on recruiting and testing participants. We would also like to thank Alejandro Pérez for his comments and suggestions on earlier versions of this manuscript.

## Notes

### Competing Interest Statement

The authors have declared no competing interest.

