## Supplementary materials for "Functional brain adaptations during speech processing in 4-month-old bilingual infants"

| SYSTEM SPECIFICATIONS | | |
|---|---|---|
| **Company and model** | NIRx NIRScout | |
| **Wavelengths** | [760, 850] | |
| **Sampling Frequency** | 15.625 Hz | |
| **Number of sources** | 8 | |
| **Number of detectors** | 12 | |
| **Number of channels** | 24 | |
| **Channel localization** | Head-based fiducial locations (10-20 system) | |
| **Sorce-Detector distances** | [20 - 30 mm] | |
| **STEPS** | **PARAMETERS** | **FUNCTION/TOOLBOX** |
| ***Data preprocessing*** | | |
| **1) Intensity to optical density** | | hmrIntensity2OD |
| **2) Motion detection and censoring** | tMask: 5; tMotion: 0.1; STDEVthresh: 20; AMPthresh: 0.2 | hmrMotionArtifact |
| **3) Motion correction** | wavelet: d4; threshold: 0.02; boundary: reflection; chsearch: moderate; nscale: extreme | *BrainWavelet Toolbox* |
| **4) Optical density to HbO and HbR** | DPF: [5.3, 4.2] | hmrOD2conc |
| **5) Low pass filter** | LPF: 0.5 Hz | hmrBandpassFilt |
| **6) Nuisance regression** | 8$^{th}$ order Legendre polynomials; mean (HbO, HbR) signal | In house script |
| ***Data analysis*** | | |
| **Detection based GLM** | glmSolveMethod: 1 (OLS); paramsBasis: [0 4 11] | hmrDeconvHRF_DriftSS |
| **FIR based GLM** | glmSolveMethod: 1 (OLS); paramsBasis: [1 1] | hmrDeconvHRF_DriftSS |
| ***Statistical analysis*** | | |
| **Two-way mixed ANOVA** | [FW, BW] - Repeated measures factor<br>[BIL, SP] - Between group factor | *JASP* |

**Supplementary Table 1.** Specifications of the fNIRS system used in this study, and details of the data preprocessing and analysis steps and parameters used in each of them. DPF: Differential pathlength factor; LPF: Low pass filter; OLS: Ordinary least squares. Functions starting with *hmr* are part of the Homer2 software package (Huppert et al., 2009; https://homer-fnirs.org). BrainWavelet Toolbox (Patel et al., 2014; http://www.brainwavelet.org/about).

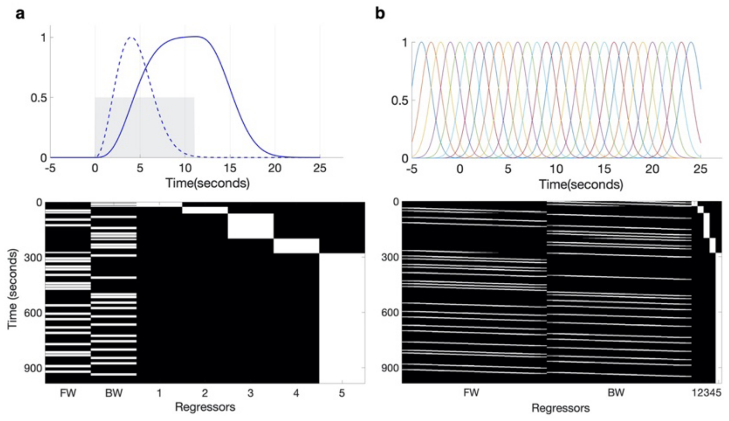


**Supplementary Fig. A.** GLM modelling of the HbO and HbR signals for each condition using regressors time-locked to stimuli. **a)** First GLM analysis with the convolution of a gamma function with peak at four seconds (dashed blue) and a square wave of eleven seconds (grey). **b)** Second GLM analysis was a finite impulse response (a.k.a., deconvolution) model using a set of gaussian basis functions

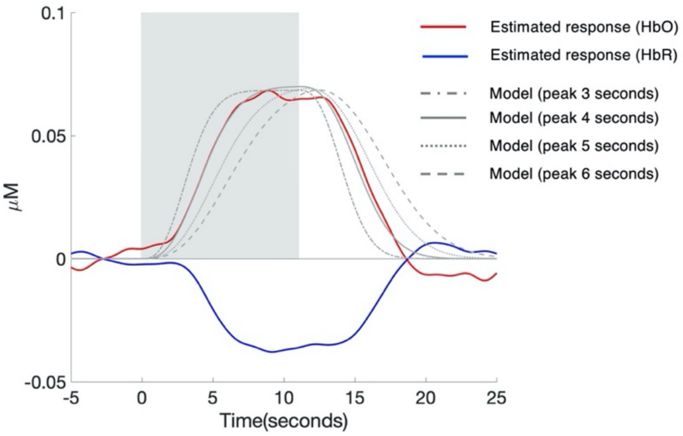


**Supplementary Fig. B.** Comparison between the estimated hemodynamic response based on the FIR deconvolution model, and the model used in the detection based GLM generated by convolving a gamma function with different peak times (i.e., 3, 4, 5 and 6 seconds) with a boxcar function of eleven seconds duration. The estimated response time course was computed by averaging a subset of channels showing activation for the experimental conditions across all participants (i.e., channels 1, 2, 3, 4, 7, 9, 13, 14, 16, 17, 18, 19, 21).

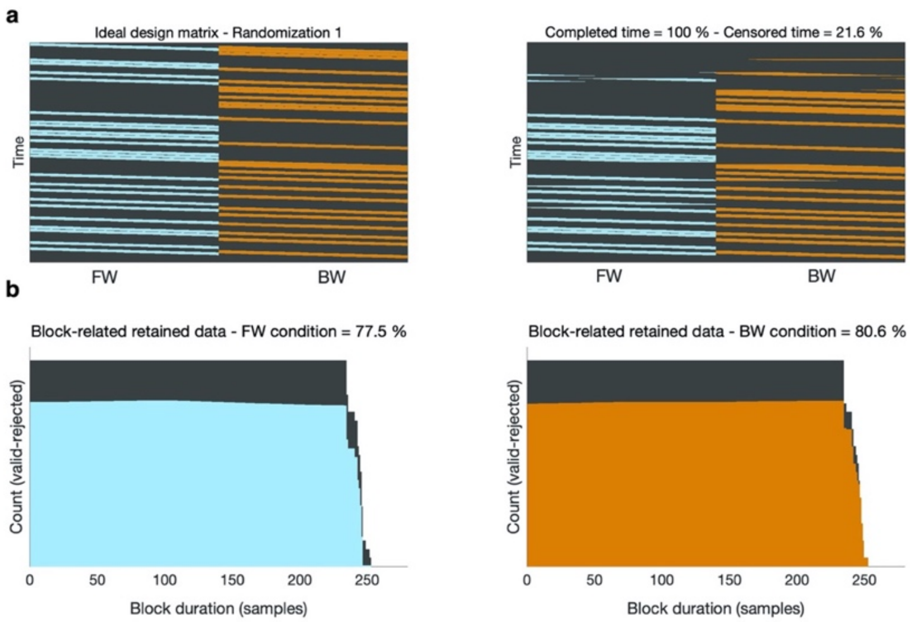


**Supplementary Fig. C.** Motion affected time points were censored for data analysis. **a)** Ideal design matrix with all-time points included for both conditions (left) and evaluation of the percentage of block-related time censored on an exemplary participant (right). **b)** In this example, and for both experimental conditions, all time-points inside each block contributed with a similar amount of data to statistical analyses.

| BIL - FW | | | | | | |
|---|---|---|---|---|---|---|
| **CH** | **Hb** | **t-value** | **p-value** | **CI** | | |
| **1** | **HbO** | -1.872 | 0.073 | -0.301 | 0.014 | |
| **2** | **HbO** | -2.797 | 0.010 | -0.422 | -0.064 | * |
| **3** | **HbO** | -1.542 | 0.136 | -0.414 | 0.059 | |
| **4** | **HbO** | -3.550 | 0.002 | -0.389 | -0.103 | * |
| **5** | **HbO** | 0.783 | 0.441 | -0.160 | 0.355 | |
| **6** | **HbO** | 0.485 | 0.632 | -0.186 | 0.300 | |
| **7** | **HbO** | 3.123 | 0.004 | 0.113 | 0.551 | * |
| **8** | **HbO** | -1.928 | 0.065 | -0.257 | 0.008 | |
| **9** | **HbO** | 1.826 | 0.080 | -0.024 | 0.394 | |
| **10** | **HbO** | 0.916 | 0.368 | -0.074 | 0.193 | |
| **11** | **HbO** | 0.521 | 0.607 | -0.115 | 0.193 | |
| **12** | **HbO** | -0.574 | 0.571 | -0.184 | 0.104 | |
| **13** | **HbO** | -3.861 | 0.001 | -0.300 | -0.091 | * |
| **14** | **HbO** | -3.722 | 0.001 | -0.453 | -0.130 | * |
| **15** | **HbO** | -2.305 | 0.030 | -0.246 | -0.014 | |
| **16** | **HbO** | -4.319 | 0.000 | -0.412 | -0.146 | * |
| **17** | **HbO** | 2.617 | 0.015 | 0.059 | 0.496 | * |
| **18** | **HbO** | 2.221 | 0.036 | 0.017 | 0.450 | |
| **19** | **HbO** | 5.248 | 0.000 | 0.300 | 0.688 | * |
| **20** | **HbO** | -0.382 | 0.706 | -0.182 | 0.125 | |
| **21** | **HbO** | 2.557 | 0.017 | 0.042 | 0.387 | * |
| **22** | **HbO** | -0.400 | 0.693 | -0.130 | 0.088 | |
| **23** | **HbO** | 0.106 | 0.916 | -0.101 | 0.112 | |
| **24** | **HbO** | -0.972 | 0.340 | -0.235 | 0.084 | |
| **1** | **HbR** | 3.051 | 0.005 | 0.027 | 0.140 | |
| **2** | **HbR** | 4.984 | 0.000 | 0.090 | 0.217 | * |
| **3** | **HbR** | 0.430 | 0.671 | -0.079 | 0.120 | * |
| **4** | **HbR** | 3.612 | 0.001 | 0.051 | 0.188 | |
| **5** | **HbR** | -0.398 | 0.694 | -0.156 | 0.106 | * |
| **6** | **HbR** | -0.695 | 0.494 | -0.182 | 0.090 | |
| **7** | **HbR** | -3.257 | 0.003 | -0.286 | -0.064 | |
| **8** | **HbR** | 2.301 | 0.030 | 0.007 | 0.136 | * |
| **9** | **HbR** | -2.288 | 0.031 | -0.174 | -0.009 | |
| **10** | **HbR** | -1.318 | 0.200 | -0.089 | 0.020 | |
| **11** | **HbR** | -1.797 | 0.084 | -0.121 | 0.008 | |
| **12** | **HbR** | -1.355 | 0.187 | -0.090 | 0.018 | |
| **13** | **HbR** | 3.491 | 0.002 | 0.046 | 0.180 | |
| **14** | **HbR** | 3.264 | 0.003 | 0.049 | 0.215 | * |
| **15** | **HbR** | 2.333 | 0.028 | 0.009 | 0.143 | * |
| **16** | **HbR** | 5.722 | 0.000 | 0.101 | 0.214 | |
| **17** | **HbR** | -2.432 | 0.023 | -0.219 | -0.018 | * |
| **18** | **HbR** | -1.621 | 0.118 | -0.208 | 0.025 | |
| **19** | **HbR** | -5.468 | 0.000 | -0.330 | -0.150 | |
| **20** | **HbR** | 0.671 | 0.508 | -0.043 | 0.084 | * |
| **21** | **HbR** | -0.869 | 0.393 | -0.131 | 0.053 | |
| **22** | **HbR** | -0.180 | 0.858 | -0.060 | 0.050 | |
| **23** | **HbR** | 0.163 | 0.872 | -0.042 | 0.049 | |
| **24** | **HbR** | 0.252 | 0.803 | -0.052 | 0.066 | |

**Supplementary Table 2.** Channelwise results of one-sample t-tests conducted in the BIL group for the FW condition (HbO and HbR). * Denotes effects that survived multiple comparisons correction.

| | | BIL - BW | | | | |
|---|---|---|---|---|---|---|
| **CH** | **Hb** | **t-value** | **p-value** | **CI** | | |
| **1** | **HbO** | -1.901 | 0.069 | -0.305 | 0.012 | |
| **2** | **HbO** | -3.280 | 0.003 | -0.409 | -0.094 | * |
| **3** | **HbO** | -0.997 | 0.328 | -0.361 | 0.125 | |
| **4** | **HbO** | -3.273 | 0.003 | -0.436 | -0.099 | * |
| **5** | **HbO** | 0.314 | 0.756 | -0.184 | 0.250 | |
| **6** | **HbO** | 1.352 | 0.188 | -0.085 | 0.408 | |
| **7** | **HbO** | 3.541 | 0.002 | 0.160 | 0.604 | * |
| **8** | **HbO** | -2.720 | 0.012 | -0.273 | -0.038 | * |
| **9** | **HbO** | 2.557 | 0.017 | 0.044 | 0.407 | * |
| **10** | **HbO** | -0.103 | 0.919 | -0.134 | 0.122 | |
| **11** | **HbO** | 0.576 | 0.570 | -0.093 | 0.166 | |
| **12** | **HbO** | 0.002 | 0.998 | -0.150 | 0.150 | |
| **13** | **HbO** | -5.639 | 0.000 | -0.352 | -0.164 | * |
| **14** | **HbO** | -4.822 | 0.000 | -0.386 | -0.155 | * |
| **15** | **HbO** | -3.286 | 0.003 | -0.376 | -0.086 | * |
| **16** | **HbO** | -5.351 | 0.000 | -0.451 | -0.200 | * |
| **17** | **HbO** | 3.388 | 0.002 | 0.104 | 0.425 | * |
| **18** | **HbO** | 2.406 | 0.024 | 0.035 | 0.455 | * |
| **19** | **HbO** | 4.554 | 0.000 | 0.226 | 0.600 | * |
| **20** | **HbO** | -0.351 | 0.729 | -0.143 | 0.101 | |
| **21** | **HbO** | 3.982 | 0.001 | 0.134 | 0.420 | * |
| **22** | **HbO** | 0.517 | 0.610 | -0.115 | 0.192 | |
| **23** | **HbO** | -0.033 | 0.974 | -0.087 | 0.084 | |
| **24** | **HbO** | -0.388 | 0.701 | -0.162 | 0.111 | |
| **1** | **HbR** | 2.148 | 0.042 | 0.003 | 0.136 | |
| **2** | **HbR** | 4.313 | 0.000 | 0.070 | 0.198 | * |
| **3** | **HbR** | 0.559 | 0.581 | -0.083 | 0.144 | |
| **4** | **HbR** | 3.479 | 0.002 | 0.048 | 0.187 | * |
| **5** | **HbR** | -0.086 | 0.932 | -0.120 | 0.110 | |
| **6** | **HbR** | -1.041 | 0.308 | -0.217 | 0.071 | |
| **7** | **HbR** | -3.412 | 0.002 | -0.305 | -0.075 | * |
| **8** | **HbR** | 2.267 | 0.032 | 0.006 | 0.130 | |
| **9** | **HbR** | -1.958 | 0.061 | -0.157 | 0.004 | |
| **10** | **HbR** | -1.809 | 0.083 | -0.100 | 0.006 | |
| **11** | **HbR** | -1.921 | 0.066 | -0.082 | 0.003 | |
| **12** | **HbR** | -0.972 | 0.341 | -0.064 | 0.023 | |
| **13** | **HbR** | 5.439 | 0.000 | 0.070 | 0.156 | * |
| **14** | **HbR** | 4.865 | 0.000 | 0.079 | 0.195 | * |
| **15** | **HbR** | 3.035 | 0.006 | 0.036 | 0.190 | * |
| **16** | **HbR** | 3.728 | 0.001 | 0.048 | 0.168 | * |
| **17** | **HbR** | -2.926 | 0.007 | -0.197 | -0.034 | * |
| **18** | **HbR** | -1.923 | 0.066 | -0.206 | 0.007 | |
| **19** | **HbR** | -4.884 | 0.000 | -0.278 | -0.113 | * |
| **20** | **HbR** | 1.173 | 0.252 | -0.026 | 0.094 | |
| **21** | **HbR** | -3.195 | 0.004 | -0.145 | -0.031 | * |
| **22** | **HbR** | -0.227 | 0.822 | -0.080 | 0.064 | |
| **23** | **HbR** | 0.331 | 0.744 | -0.033 | 0.045 | |
| **24** | **HbR** | 0.818 | 0.421 | -0.041 | 0.094 | |

**Supplementary Table 3.** Channelwise results of one-sample t-tests conducted in the BIL group for the BW condition (HbO and HbR). * Denotes effects that survived multiple comparisons correction.

| | | SP - FW | | | | |
|---|---|---|---|---|---|---|
| **CH** | **Hb** | **t-value** | **p-value** | **CI** | | |
| **1** | **HbO** | -2.953 | 0.006 | -0.209 | -0.038 | |
| **2** | **HbO** | -3.614 | 0.001 | -0.302 | -0.084 | * |
| **3** | **HbO** | -0.683 | 0.500 | -0.245 | 0.122 | |
| **4** | **HbO** | -2.801 | 0.009 | -0.327 | -0.051 | |
| **5** | **HbO** | 1.040 | 0.307 | -0.082 | 0.252 | |
| **6** | **HbO** | -0.717 | 0.479 | -0.196 | 0.094 | |
| **7** | **HbO** | 1.916 | 0.065 | -0.008 | 0.258 | |
| **8** | **HbO** | -1.601 | 0.120 | -0.184 | 0.022 | |
| **9** | **HbO** | -0.293 | 0.771 | -0.159 | 0.119 | |
| **10** | **HbO** | -0.864 | 0.394 | -0.145 | 0.059 | |
| **11** | **HbO** | 0.402 | 0.690 | -0.068 | 0.102 | |
| **12** | **HbO** | -1.208 | 0.236 | -0.167 | 0.043 | |
| **13** | **HbO** | -0.176 | 0.862 | -0.090 | 0.075 | |
| **14** | **HbO** | -0.946 | 0.352 | -0.175 | 0.064 | |
| **15** | **HbO** | 0.482 | 0.633 | -0.127 | 0.206 | |
| **16** | **HbO** | 0.742 | 0.464 | -0.057 | 0.123 | |
| **17** | **HbO** | 1.921 | 0.064 | -0.008 | 0.250 | |
| **18** | **HbO** | 2.412 | 0.022 | 0.034 | 0.405 | |
| **19** | **HbO** | 2.092 | 0.045 | 0.003 | 0.257 | |
| **20** | **HbO** | 0.558 | 0.581 | -0.097 | 0.170 | |
| **21** | **HbO** | 1.057 | 0.299 | -0.053 | 0.168 | |
| **22** | **HbO** | 0.227 | 0.822 | -0.107 | 0.134 | |
| **23** | **HbO** | 0.117 | 0.907 | -0.089 | 0.100 | |
| **24** | **HbO** | 0.062 | 0.951 | -0.120 | 0.127 | |
| **1** | **HbR** | 2.197 | 0.036 | 0.004 | 0.101 | |
| **2** | **HbR** | 2.309 | 0.028 | 0.008 | 0.127 | |
| **3** | **HbR** | 1.199 | 0.240 | -0.037 | 0.142 | |
| **4** | **HbR** | 1.457 | 0.156 | -0.018 | 0.109 | |
| **5** | **HbR** | 0.663 | 0.513 | -0.053 | 0.105 | |
| **6** | **HbR** | 0.906 | 0.372 | -0.043 | 0.112 | |
| **7** | **HbR** | -2.403 | 0.023 | -0.145 | -0.012 | |
| **8** | **HbR** | 0.530 | 0.600 | -0.050 | 0.085 | |
| **9** | **HbR** | 1.580 | 0.125 | -0.013 | 0.104 | |
| **10** | **HbR** | -0.707 | 0.485 | -0.077 | 0.037 | |
| **11** | **HbR** | 0.926 | 0.362 | -0.019 | 0.051 | |
| **12** | **HbR** | 0.909 | 0.371 | -0.035 | 0.090 | |
| **13** | **HbR** | 1.784 | 0.085 | -0.006 | 0.090 | |
| **14** | **HbR** | 1.943 | 0.061 | -0.003 | 0.125 | |
| **15** | **HbR** | 0.257 | 0.799 | -0.082 | 0.106 | |
| **16** | **HbR** | 1.319 | 0.197 | -0.016 | 0.076 | |
| **17** | **HbR** | -3.884 | 0.001 | -0.133 | -0.041 | * |
| **18** | **HbR** | -2.666 | 0.012 | -0.181 | -0.024 | |
| **19** | **HbR** | -1.741 | 0.092 | -0.094 | 0.007 | |
| **20** | **HbR** | -0.523 | 0.605 | -0.071 | 0.042 | |
| **21** | **HbR** | -2.021 | 0.052 | -0.123 | 0.001 | |
| **22** | **HbR** | -1.674 | 0.105 | -0.108 | 0.011 | |
| **23** | **HbR** | -1.285 | 0.209 | -0.078 | 0.018 | |
| **24** | **HbR** | -1.412 | 0.168 | -0.107 | 0.020 | |

**Supplementary Table 4.** Channelwise results of one-sample t-tests conducted in the SP group for the FW condition (HbO and HbR). * Denotes effects that survived multiple comparisons correction.

| | | SP - BW | | | | |
|---|---|---|---|---|---|---|
| **CH** | **Hb** | **t-value** | **p-value** | **CI** | | |
| **1** | **HbO** | -2.532 | 0.017 | -0.236 | -0.025 | |
| **2** | **HbO** | -3.359 | 0.002 | -0.346 | -0.084 | * |
| **3** | **HbO** | -0.801 | 0.429 | -0.276 | 0.121 | |
| **4** | **HbO** | -1.508 | 0.142 | -0.219 | 0.033 | |
| **5** | **HbO** | 0.802 | 0.429 | -0.109 | 0.249 | |
| **6** | **HbO** | 0.383 | 0.705 | -0.132 | 0.193 | |
| **7** | **HbO** | 1.955 | 0.060 | -0.006 | 0.284 | |
| **8** | **HbO** | -1.181 | 0.247 | -0.158 | 0.042 | |
| **9** | **HbO** | 0.036 | 0.972 | -0.130 | 0.135 | |
| **10** | **HbO** | -1.564 | 0.128 | -0.182 | 0.024 | |
| **11** | **HbO** | 0.425 | 0.674 | -0.081 | 0.123 | |
| **12** | **HbO** | -2.068 | 0.047 | -0.243 | -0.002 | |
| **13** | **HbO** | -3.225 | 0.003 | -0.135 | -0.030 | * |
| **14** | **HbO** | -2.659 | 0.012 | -0.225 | -0.029 | * |
| **15** | **HbO** | 1.196 | 0.241 | -0.046 | 0.174 | |
| **16** | **HbO** | -1.433 | 0.162 | -0.185 | 0.032 | |
| **17** | **HbO** | 4.259 | 0.000 | 0.104 | 0.297 | * |
| **18** | **HbO** | 2.909 | 0.007 | 0.088 | 0.502 | * |
| **19** | **HbO** | 3.327 | 0.002 | 0.081 | 0.337 | * |
| **20** | **HbO** | -0.220 | 0.827 | -0.135 | 0.109 | |
| **21** | **HbO** | 1.914 | 0.065 | -0.008 | 0.236 | |
| **22** | **HbO** | 0.439 | 0.664 | -0.089 | 0.138 | |
| **23** | **HbO** | -0.689 | 0.496 | -0.129 | 0.064 | |
| **24** | **HbO** | -0.808 | 0.426 | -0.220 | 0.095 | |
| **1** | **HbR** | 2.458 | 0.020 | 0.009 | 0.099 | * |
| **2** | **HbR** | 3.139 | 0.004 | 0.028 | 0.132 | * |
| **3** | **HbR** | 1.248 | 0.222 | -0.035 | 0.145 | |
| **4** | **HbR** | 1.678 | 0.104 | -0.009 | 0.096 | |
| **5** | **HbR** | 0.078 | 0.938 | -0.085 | 0.092 | |
| **6** | **HbR** | 0.985 | 0.333 | -0.030 | 0.086 | |
| **7** | **HbR** | -1.424 | 0.165 | -0.114 | 0.020 | |
| **8** | **HbR** | 1.400 | 0.172 | -0.011 | 0.058 | |
| **9** | **HbR** | -0.506 | 0.617 | -0.070 | 0.042 | |
| **10** | **HbR** | 0.387 | 0.702 | -0.044 | 0.065 | |
| **11** | **HbR** | 0.315 | 0.755 | -0.036 | 0.049 | |
| **12** | **HbR** | 0.856 | 0.399 | -0.029 | 0.070 | |
| **13** | **HbR** | 3.185 | 0.003 | 0.020 | 0.092 | * |
| **14** | **HbR** | 3.284 | 0.003 | 0.030 | 0.130 | * |
| **15** | **HbR** | 1.215 | 0.234 | -0.027 | 0.106 | |
| **16** | **HbR** | 4.170 | 0.000 | 0.050 | 0.146 | * |
| **17** | **HbR** | -3.694 | 0.001 | -0.145 | -0.042 | * |
| **18** | **HbR** | -1.757 | 0.089 | -0.171 | 0.013 | |
| **19** | **HbR** | -3.722 | 0.001 | -0.166 | -0.048 | * |
| **20** | **HbR** | -0.222 | 0.826 | -0.057 | 0.046 | |
| **21** | **HbR** | -3.003 | 0.005 | -0.152 | -0.029 | * |
| **22** | **HbR** | -2.557 | 0.016 | -0.120 | -0.013 | * |
| **23** | **HbR** | -2.561 | 0.016 | -0.097 | -0.011 | * |
| **24** | **HbR** | -1.373 | 0.180 | -0.104 | 0.020 | |

**Supplementary Table 5.** Channelwise results of one-sample t-tests conducted in the SP group for the BW condition (HbO and HbR). * Denotes effects that survived multiple comparisons correction.

| | | Condition (FW, BW) | | |
|---|---|---|---|---|
| **CH** | **Hb** | **F-value** | **p-value** | **Partial $\eta^2$** |
| **1** | **HbO** | 0.018 | 0.894 | 0.000 |
| **2** | **HbO** | 0.104 | 0.748 | 0.002 |
| **3** | **HbO** | 0.163 | 0.688 | 0.003 |
| **4** | **HbO** | 0.619 | 0.435 | 0.011 |
| **5** | **HbO** | 0.590 | 0.446 | 0.011 |
| **6** | **HbO** | 3.698 | 0.060 | 0.063 |
| **7** | **HbO** | 0.620 | 0.435 | 0.011 |
| **8** | **HbO** | 0.008 | 0.931 | 0.000 |
| **9** | **HbO** | 0.293 | 0.591 | 0.005 |
| **10** | **HbO** | 1.302 | 0.259 | 0.023 |
| **11** | **HbO** | 0.000 | 0.985 | 0.000 |
| **12** | **HbO** | 0.040 | 0.842 | 0.001 |
| **13** | **HbO** | 4.453 | 0.039 | 0.075 |
| **14** | **HbO** | 0.350 | 0.556 | 0.006 |
| **15** | **HbO** | 0.651 | 0.423 | 0.012 |
| **16** | **HbO** | 2.820 | 0.099 | 0.049 |
| **17** | **HbO** | 0.730 | 0.397 | 0.013 |
| **18** | **HbO** | 0.627 | 0.432 | 0.011 |
| **19** | **HbO** | 0.001 | 0.979 | 0.000 |
| **20** | **HbO** | 0.363 | 0.549 | 0.007 |
| **21** | **HbO** | 2.288 | 0.136 | 0.040 |
| **22** | **HbO** | 0.780 | 0.381 | 0.014 |
| **23** | **HbO** | 0.407 | 0.526 | 0.007 |
| **24** | **HbO** | 0.025 | 0.875 | 0.000 |
| **1** | **HbR** | 0.153 | 0.697 | 0.003 |
| **2** | **HbR** | 0.030 | 0.863 | 0.001 |
| **3** | **HbR** | 0.058 | 0.810 | 0.001 |
| **4** | **HbR** | 0.011 | 0.918 | 0.000 |
| **5** | **HbR** | 0.001 | 0.970 | 0.000 |
| **6** | **HbR** | 0.468 | 0.497 | 0.008 |
| **7** | **HbR** | 0.192 | 0.663 | 0.003 |
| **8** | **HbR** | 0.004 | 0.950 | 0.000 |
| **9** | **HbR** | 0.963 | 0.331 | 0.017 |
| **10** | **HbR** | 0.250 | 0.619 | 0.005 |
| **11** | **HbR** | 0.053 | 0.819 | 0.001 |
| **12** | **HbR** | 0.042 | 0.838 | 0.001 |
| **13** | **HbR** | 0.129 | 0.721 | 0.002 |
| **14** | **HbR** | 0.371 | 0.545 | 0.007 |
| **15** | **HbR** | 1.728 | 0.194 | 0.030 |
| **16** | **HbR** | 0.334 | 0.566 | 0.006 |
| **17** | **HbR** | 0.008 | 0.927 | 0.000 |
| **18** | **HbR** | 0.086 | 0.770 | 0.002 |
| **19** | **HbR** | 0.336 | 0.564 | 0.006 |
| **20** | **HbR** | 0.285 | 0.596 | 0.005 |
| **21** | **HbR** | 3.069 | 0.085 | 0.053 |
| **22** | **HbR** | 0.288 | 0.594 | 0.005 |
| **23** | **HbR** | 0.350 | 0.557 | 0.006 |
| **24** | **HbR** | 0.262 | 0.611 | 0.005 |

**Supplementary Table 6.** Channelwise results of the two-way mixed ANOVA for the main effect of Condition (HbO and HbR). * Denotes effects that survived multiple comparisons correction.

| | | Language (BIL, SP) | | | |
|---|---|---|---|---|---|
| **CH** | **Hb** | **F-value** | **p-value** | **Partial $\eta^2$** | |
| **1** | **HbO** | 0.053 | 0.819 | 0.001 | |
| **2** | **HbO** | 0.250 | 0.619 | 0.005 | |
| **3** | **HbO** | 0.319 | 0.575 | 0.006 | |
| **4** | **HbO** | 1.781 | 0.188 | 0.031 | |
| **5** | **HbO** | 0.009 | 0.927 | 0.000 | |
| **6** | **HbO** | 0.882 | 0.352 | 0.016 | |
| **7** | **HbO** | 3.738 | 0.058 | 0.064 | |
| **8** | **HbO** | 1.184 | 0.281 | 0.021 | |
| **9** | **HbO** | 4.848 | 0.032 | 0.081 | |
| **10** | **HbO** | 1.742 | 0.192 | 0.031 | |
| **11** | **HbO** | 0.077 | 0.783 | 0.001 | |
| **12** | **HbO** | 0.950 | 0.334 | 0.017 | |
| **13** | **HbO** | 14.696 | <0.001 | 0.211 | * |
| **14** | **HbO** | 6.609 | 0.013 | 0.107 | |
| **15** | **HbO** | 7.839 | 0.007 | 0.125 | * |
| **16** | **HbO** | 19.610 | <0.001 | 0.263 | * |
| **17** | **HbO** | 1.281 | 0.263 | 0.023 | |
| **18** | **HbO** | 0.018 | 0.892 | 0.000 | |
| **19** | **HbO** | 7.806 | 0.007 | 0.124 | * |
| **20** | **HbO** | 0.181 | 0.672 | 0.003 | |
| **21** | **HbO** | 3.530 | 0.066 | 0.060 | |
| **22** | **HbO** | 0.018 | 0.894 | 0.000 | |
| **23** | **HbO** | 0.076 | 0.783 | 0.001 | |
| **24** | **HbO** | 0.061 | 0.806 | 0.001 | |
| **1** | **HbR** | 0.481 | 0.491 | 0.009 | |
| **2** | **HbR** | 3.928 | 0.053 | 0.067 | |
| **3** | **HbR** | 0.203 | 0.654 | 0.004 | |
| **4** | **HbR** | 3.669 | 0.061 | 0.063 | |
| **5** | **HbR** | 0.196 | 0.660 | 0.004 | |
| **6** | **HbR** | 1.778 | 0.188 | 0.031 | |
| **7** | **HbR** | 4.186 | 0.046 | 0.071 | |
| **8** | **HbR** | 1.998 | 0.163 | 0.035 | |
| **9** | **HbR** | 5.729 | 0.020 | 0.094 | |
| **10** | **HbR** | 1.152 | 0.288 | 0.021 | |
| **11** | **HbR** | 4.613 | 0.036 | 0.077 | |
| **12** | **HbR** | 2.720 | 0.105 | 0.047 | |
| **13** | **HbR** | 5.544 | 0.022 | 0.092 | |
| **14** | **HbR** | 2.650 | 0.109 | 0.046 | |
| **15** | **HbR** | 2.042 | 0.159 | 0.036 | |
| **16** | **HbR** | 4.650 | 0.035 | 0.078 | |
| **17** | **HbR** | 0.349 | 0.557 | 0.006 | |
| **18** | **HbR** | 0.006 | 0.938 | 0.000 | |
| **19** | **HbR** | 9.871 | 0.003 | 0.152 | |
| **20** | **HbR** | 1.217 | 0.275 | 0.022 | |
| **21** | **HbR** | 0.086 | 0.771 | 0.002 | |
| **22** | **HbR** | 1.977 | 0.165 | 0.035 | |
| **23** | **HbR** | 3.575 | 0.064 | 0.061 | |
| **24** | **HbR** | 2.411 | 0.126 | 0.042 | |

**Supplementary Table 7.** Channelwise results of the two-way mixed ANOVA for the main effect of Language (HbO and HbR). * Denotes effects that survived multiple comparisons correction.

| CH | Hb | Language*Condition | | | |
| --- | --- | --- | --- | --- | --- |
| | | F-value | p-value | Partial $\eta^2$ | |
| **1** | **HbO** | 0.003 | 0.957 | 0.000 | |
| **2** | **HbO** | 0.022 | 0.884 | 0.000 | |
| **3** | **HbO** | 0.516 | 0.476 | 0.009 | |
| **4** | **HbO** | 1.535 | 0.221 | 0.027 | |
| **5** | **HbO** | 0.233 | 0.631 | 0.004 | |
| **6** | **HbO** | 0.058 | 0.811 | 0.001 | |
| **7** | **HbO** | 0.197 | 0.659 | 0.004 | |
| **8** | **HbO** | 0.375 | 0.543 | 0.007 | |
| **9** | **HbO** | 0.025 | 0.876 | 0.000 | |
| **10** | **HbO** | 0.114 | 0.736 | 0.002 | |
| **11** | **HbO** | 0.007 | 0.935 | 0.000 | |
| **12** | **HbO** | 1.033 | 0.314 | 0.018 | |
| **13** | **HbO** | 0.043 | 0.837 | 0.001 | |
| **14** | **HbO** | 1.175 | 0.283 | 0.021 | |
| **15** | **HbO** | 1.789 | 0.186 | 0.032 | |
| **16** | **HbO** | 0.459 | 0.501 | 0.008 | |
| **17** | **HbO** | 1.429 | 0.237 | 0.025 | |
| **18** | **HbO** | 0.331 | 0.567 | 0.006 | |
| **19** | **HbO** | 4.281 | 0.043 | 0.072 | |
| **20** | **HbO** | 0.676 | 0.414 | 0.012 | |
| **21** | **HbO** | 0.006 | 0.939 | 0.000 | |
| **22** | **HbO** | 0.371 | 0.545 | 0.007 | |
| **23** | **HbO** | 0.197 | 0.659 | 0.004 | |
| **24** | **HbO** | 1.247 | 0.269 | 0.022 | |
| **1** | **HbR** | 0.255 | 0.616 | 0.005 | |
| **2** | **HbR** | 0.561 | 0.457 | 0.010 | |
| **3** | **HbR** | 0.021 | 0.886 | 0.000 | |
| **4** | **HbR** | 0.000 | 0.994 | 0.000 | |
| **5** | **HbR** | 0.824 | 0.368 | 0.015 | |
| **6** | **HbR** | 0.177 | 0.676 | 0.003 | |
| **7** | **HbR** | 1.451 | 0.233 | 0.026 | |
| **8** | **HbR** | 0.061 | 0.805 | 0.001 | |
| **9** | **HbR** | 2.641 | 0.110 | 0.046 | |
| **10** | **HbR** | 1.287 | 0.262 | 0.023 | |
| **11** | **HbR** | 0.650 | 0.423 | 0.012 | |
| **12** | **HbR** | 0.318 | 0.575 | 0.006 | |
| **13** | **HbR** | 0.126 | 0.724 | 0.002 | |
| **14** | **HbR** | 0.107 | 0.745 | 0.002 | |
| **15** | **HbR** | 0.038 | 0.846 | 0.001 | |
| **16** | **HbR** | 11.899 | 0.001 | 0.178 | * |
| **17** | **HbR** | 0.067 | 0.797 | 0.001 | |
| **18** | **HbR** | 0.374 | 0.544 | 0.007 | |
| **19** | **HbR** | 10.551 | 0.002 | 0.161 | * |
| **20** | **HbR** | 0.012 | 0.915 | 0.000 | |
| **21** | **HbR** | 0.200 | 0.657 | 0.004 | |
| **22** | **HbR** | 0.142 | 0.707 | 0.003 | |
| **23** | **HbR** | 0.544 | 0.464 | 0.010 | |
| **24** | **HbR** | 0.184 | 0.669 | 0.003 | |

**Supplementary Table 8.** Channelwise results of the two-way mixed ANOVA for the Language*Condition interaction (HbO and HbR). * Denotes effects that survived multiple comparisons correction.

**Post-hoc tests (Ch 16, HbO) – Language*Condition**

| | | Mean Difference | SE | t | $p_{tukey}$ |
|---|---|---|---|---|---|
| BIL, FW | SP, FW | -0.312 | 0.078 | -3.974 | < .001 |
| | BIL, BW | 0.046 | 0.068 | 0.680 | 0.904 |
| | SP, BW | -0.203 | 0.078 | -2.582 | 0.054 |
| SP, FW | BIL, BW | 0.358 | 0.078 | 4.565 | < .001 |
| | SP, BW | 0.109 | 0.063 | 1.745 | 0.311 |
| BIL, BW | SP, BW | -0.249 | 0.078 | -3.174 | 0.011 |

**Post-hoc tests (Ch 16, HbR) – Language*Condition**

| | | Mean Difference | SE | t | $p_{tukey}$ |
|---|---|---|---|---|---|
| BIL, FW | SP, FW | 0.127 | 0.036 | 3.528 | 0.004 |
| | BIL, BW | 0.049 | 0.025 | 1.947 | 0.221 |
| | SP, BW | 0.059 | 0.036 | 1.629 | 0.368 |
| SP, FW | BIL, BW | -0.078 | 0.036 | -2.174 | 0.139 |
| | SP, BW | -0.069 | 0.023 | -2.981 | 0.022 |
| BIL, BW | SP, BW | 0.010 | 0.036 | 0.276 | 0.993 |

**Supplementary Table 9.** Post-hoc tests on Language*Condition interaction in channel 16 for HbO (top) and HbR (bottom).

**Post-hoc tests (Ch 19, HbO) – Language*Condition**

| | | Mean Difference | SE | t | $p_{tukey}$ |
|---|---|---|---|---|---|
| BIL, FW | SP, FW | 0.364 | 0.109 | 3.345 | 0.007 |
| | BIL, BW | 0.081 | 0.057 | 1.420 | 0.492 |
| | SP, BW | 0.285 | 0.109 | 2.622 | 0.051 |
| SP, FW | BIL, BW | -0.283 | 0.109 | -2.604 | 0.053 |
| | SP, BW | -0.079 | 0.052 | -1.513 | 0.437 |
| BIL, BW | SP, BW | 0.204 | 0.109 | 1.881 | 0.245 |

**Post-hoc tests (Ch 19, HbR) – Language*Condition**

| | | Mean Difference | SE | t | $p_{tukey}$ |
|---|---|---|---|---|---|
| BIL, FW | SP, FW | -0.197 | 0.048 | -4.073 | < .001 |
| | BIL, BW | -0.045 | 0.025 | -1.809 | 0.280 |
| | SP, BW | -0.133 | 0.048 | -2.746 | 0.038 |
| SP, FW | BIL, BW | 0.152 | 0.048 | 3.148 | 0.013 |
| | SP, BW | 0.064 | 0.023 | 2.834 | 0.032 |
| BIL, BW | SP, BW | -0.088 | 0.048 | -1.821 | 0.272 |

**Supplementary Table 10.** Post-hoc tests on Language*Condition interaction in channel 19 for HbO (top) and HbR (bottom).

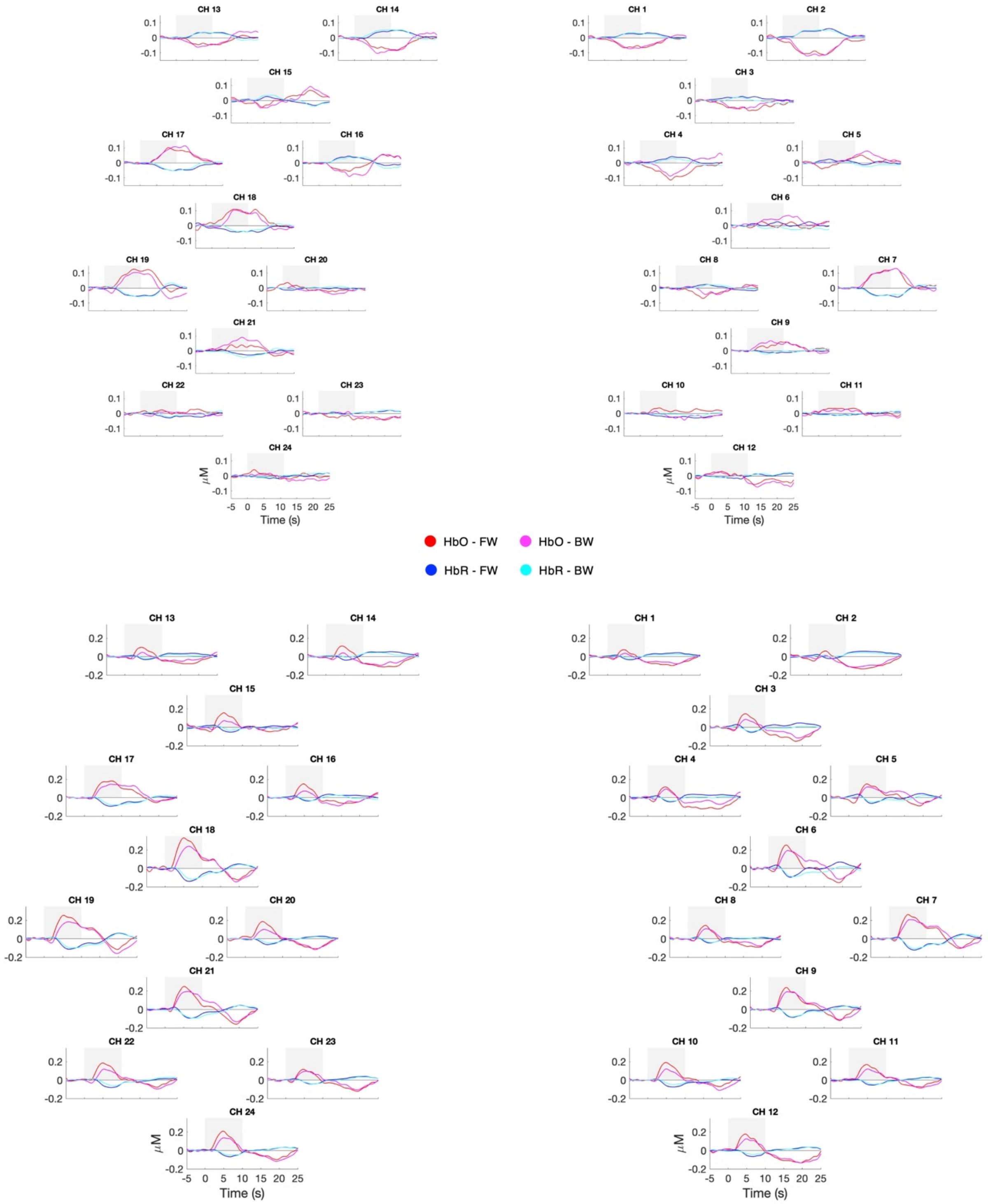


**Supplementary Fig. D.** Estimated group-level hemodynamic responses at each channel location and for each experimental condition for a data preprocessing pipeline including (top panel) and not including (bottom panel) global signal regression. Time zero indicates stimulus onset.
